# A Novel Foraging Task Reveals Cognitive and Motor Processes Underlying Behavioral Flexibility

**DOI:** 10.1101/2025.09.09.675194

**Authors:** Maud Schaffhauser, Tom Orjollet-Lacomme, Kenza Amroune, Thomas Morvan, Aurélien Fortoul, Mathias Lechelon, David Robbe

## Abstract

Adaptive behavior depends on a variety of brain functions, such as learning, decision-making, spatial navigation and motor control, which have been studied using two main strategies. Trial-based tasks allow their mechanistic dissection but tend to generate highly stereotypical behavior, whereas open-field investigations capture naturalistic dynamics with less experimental control. To leverage the strengths of both approaches, we developed a behavioral framework which recreates dilemmas faced by animals during patch foraging. In the Tower Foraging Park (TFP), mice harvest rewards along square towers (patches) by making quarter-turns around them in a single direction (exploit) and alternating between towers (explore) as patches eventually deplete. Within a couple of sessions, naïve mice performed quarter-turns in the rewarded direction with increasing vigor and reduced variability, and switched towers after short exploitation bouts. When the harvest direction was reversed, mice rapidly adapted their turning direction, with quarter-turn trajectory variability and speed becoming decoupled. Mice subjected to daily reversals adapted progressively faster, revealing meta-learning. When the next rewarding tower became harder to locate, all trained mice increased exploitation duration, although metalearners outperformed animals trained under stable contingencies. Altogether, the TFP produced behavior consistent with foraging theory and revealed new processes facilitating flexible foraging: meta-learning and the decoupling of movement variability and speed. Moreover, because the TFP accommodates diverse protocol variants, adheres to FAIR principles, and is fully compatible with modern neurophysiological techniques, it provides a promising platform for mechanistic investigations of brain functions underlying adaptive behavior while maintaining ethological validity.

## Introduction

One overarching aim of neuroscience is to understand how humans and other animals learn and flexibly adapt their behavior to satisfy needs allowing them to strive and survive. It is largely accepted that adaptive behavior relies on separate functions, such as spatial navigation, motor control, memory or sensory-guided decision-making, or procedural learning, mapping onto distinct brain regions (Shadmehr and Krakauer 2008). To understand these brain functions and their neuronal underpinnings, a widely-used strategy is to train humans and other animals on trial-based tasks that segment behavior into perceptual, cognitive, and motor epochs. This strategy reflects the necessity of simplifying the conditions under which brain functions are investigated (Chirimuuta 2024) and has led to major insights into the understanding of perception, cognition, and motor control (e.g., Georgopoulos et al. 1986; Britten et al. 1992; Basso and Wurtz 1997; Romo et al. 1999; Brunton et al. 2013). But it also faces practical challenges, especially when applied to non-humans animals. Indeed, for trial-based tasks to yield statistically robust results, animals must perform numerous trials for small rewards delivered if they follow the task’s rules (e.g., move left if stimulus A appears) and respect their epoch-based temporal structure (e.g., wait for stimulus presentation and deliberation period before responding). Yet animals are driven to secure rewards as quickly as possible (Carland et al. 2019) and spontaneously, perception, cognition, and action unfold in parallel rather than serially (Pfeifer and Bongard 2006; Cisek and Kalaska 2010; Cisek and Pastor-Bernier 2014; Wystrach 2021). Extensive training is therefore needed to counter their impulsivity and to enforce compliance with the abstract constraints of trial-based tasks. This challenge is particularly pronounced for natural foragers like rodents, especially when they are head-fixed to enable neuronal recordings (The International Brain Laboratory et al. 2021). Moreover, it has been increasingly recognized that rodents develop stereotyped uninstructed motor patterns (rituals) in such experimental contexts (Stringer et al. 2019; Musall et al. 2019; Hasnain et al. 2025; Richelle and Lejeune 1980; Killeen and Fetterman 1988), sometimes bypassing the brain functions that the experimenters aimed to study (Euston and McNaughton 2006; Cowen and McNaughton 2007; Gouvea et al. 2014; Kawai et al. 2015; Rueda-Orozco and Robbe 2015; Safaie et al. 2020; Robbe 2023). Such experimental outcome is at odds with the original scientific objective: to understand the capacity of animals to exhibit adaptive and flexible behaviors. Efforts have been made to analytically separate the neuronal correlates of movements and cognition in trial-based tasks (Hasnain et al. 2025). Still, the highly constrained structure of these tasks raises epistemologic concerns (Burgat 2010; Despret 2015; Gomez-Marin and Ghazanfar 2019; Robbe 2023; Chirimuuta 2024) and limits the understanding of the behavioral and neuronal processes that enable animals to flexibly adjust their behavior both to their dynamic needs and to environmental changes.

An alternative experimental strategy has been to take advantage of rats and mice sustained and spontaneous exploration. Historically, this approach has been successful in the study of spatial representation (O’Keefe 1979) or motor sequences (Aldridge and Berridge 1998; Berridge et al. 1987; Fentress and Stilwell 1973). Combined with recent advances in movement quantification (e.g., Mathis et al. 2018) and its unsupervised decomposition in discrete actions (e.g., Wiltschko et al. 2020), it has increasingly been used to study neuronal correlates of so-called self-paced or innate movements and social interactions (De Chaumont et al. 2012; Markowitz et al. 2018). One obvious advantage of this approach is that it does not require training and capitalize on the animal’s natural behavioral repertoire. A limitation, however, is that it may be difficult to obtain enough repetitions of complex behavioral situations to investigate higher-level functions such as sensory-guided decision-making, motor skill learning, and memory (but see Sarel et al. 2022; Markowitz et al. 2023; Sridhar et al. 2024; Ray et al. 2025). In addition, if internal and external constraints shaping spontaneous exploration and social interactions are not manipulated, the observed behavioral and neuronal dynamics can be informative but remain limited to reveal underlying mechanisms (Krakauer et al. 2017).

In summary, trial-based tasks provide analytical power at the cost of ecological validity, whereas open-field exploration captures the richness of spontaneous behavioral activity but often remains descriptive. These approaches highlight the diversity of strategies for studying behavior and its underlying brain functions, but one may wonder how to combine them to harness their strengths. We propose that patch foraging offers such an intersection. Patch foraging consists of searching for and consuming resources that are concentrated in discrete locations (Stephens and Krebs 1986; Dennis et al. 2021; Barack 2024). During patch foraging, hungry or thirsty animals must decide which patch to harvest (exploit), and after this initial commitment, must decide when to leave the patch to search for a new one (explore), hopefully in a better state than the one they left. This decision dilemma is referred to as the exploration-exploitation tradeoff. Patch foraging provides several interesting features to study adaptive behavior in a quantitative manner. First, foraging consists of a continuous cycle of decisions and actions that can be easily parsed into exploitative and exploratory phases, eliminating the need for artificial segmentation of behavior while providing enough repetitions to yield statistically robust results. Second, the decisions involved do not require associations between abstract stimuli or actions and reward delivery, avoiding lengthy training to condition animals. Third, patch foraging allows the investigation motor control, both during exploitation within a patch and during exploration between patches. Fourth, because patches eventually deplete, animals can not simply perform the same action to obtain rewards, creating a natural balance between structure and flexibility. Fifth, even if there is a theoretical optimal solution to the patch foraging problem (Charnov 1976), animals typically deviate from optimality (e.g., Kilpatrick et al. 2021). Indeed, the exploration-exploitation tradeoff is known to be sensitive to internal and external constraints (e.g., motivational state, environmental knowledge, effort) that can be manipulated, providing a straightforward approach to probe the behavioral and neuronal mechanisms underlying adaptive behavior.

Within the constraints of a laboratory, we developed a novel patch-foraging paradigm that we named the Towers Foraging Park (TFP). We tested two large groups of mice using distinct foraging protocols. The results show that mice rapidly learned to forage in this type of task, exhibiting adaptive and flexible exploration-exploitation tradeoffs, procedural learning, and reward-related modulations of movement kinematics. The specific protocols allowed us to modify the foraging rules, revealing two mechanisms that promote flexible foraging: a cognitive one (metalearning) and a motor one (the uncoupling of movement speed and variability). Overall, these findings highlight the value of ethologically grounded experiments for studying adaptive behavior and pave the way for future investigations of its neuronal underpinnings.

## Results

To emulate the dilemmas faced by animals during patch foraging we created an experimental apparatus consisting of a custom-made square arena in which four smaller square towers are positioned at the center of each quadrant (northwest, northeast, southwest, and southeast tower, Figure 1AB). The four walls of each tower (north, east, south, and west) are equipped with a small tube located at its center (Figure 1B right). Each tube (referred to as a water port in the rest of the manuscript) is connected to a water reservoir housed within the tower to deliver single water drops. The arena floor is transparent, allowing video tracking of the mice’s position using an infrared camera positioned beneath the apparatus. The experiments are conducted in complete darkness.

**Figure 1.**
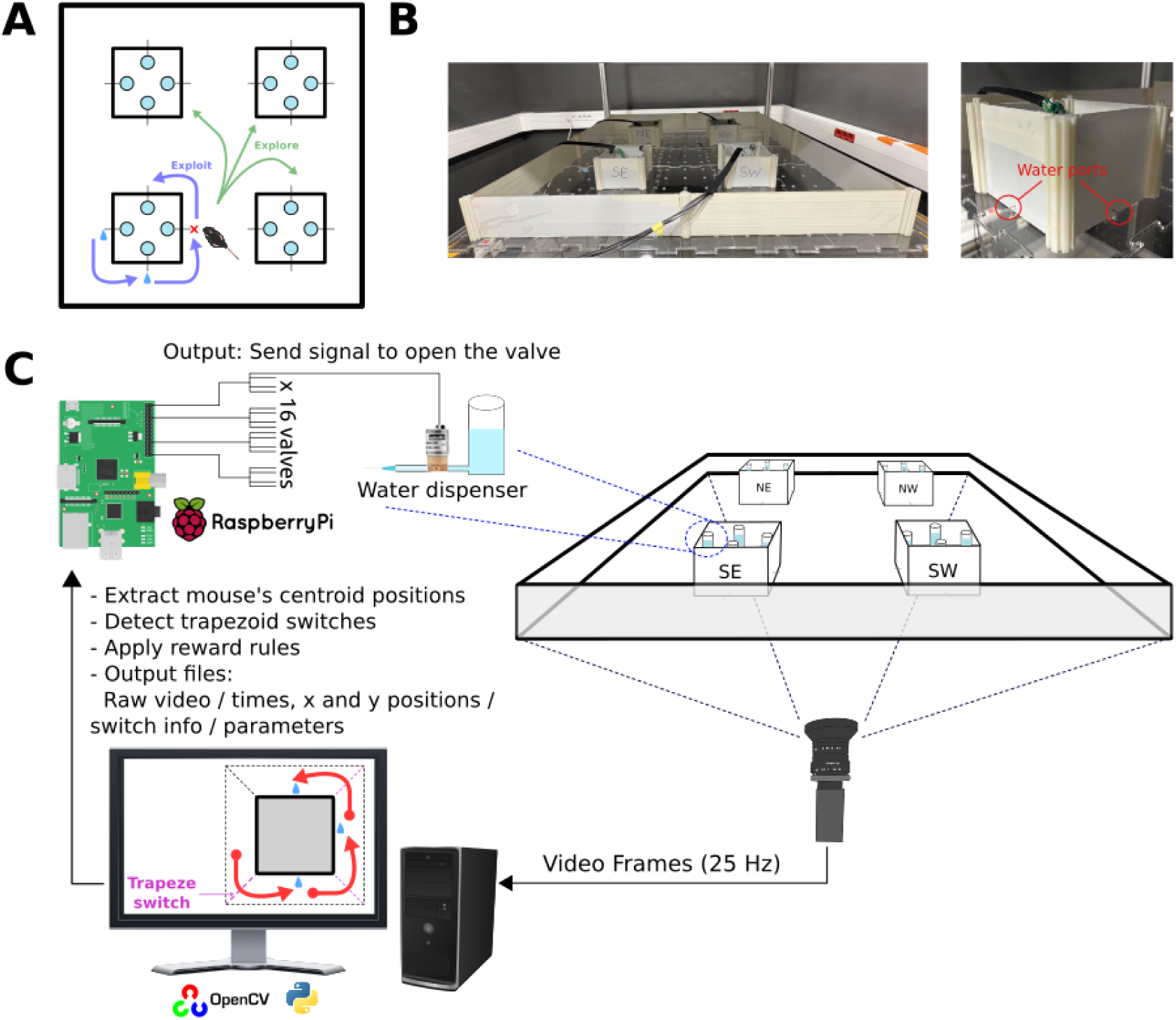
Schematic and control principle of the Towers Foraging Park. **A**) Schematic representation of the task structure. Mice perform repetitive reward-oriented quarter-turns around a tower to obtain drops of water (exploitation), and transitions to other towers (exploration). **B**) Photographs of the full arena (left) and a close-up of a tower showing the position of two water ports (red circles, right). **C**) Overview of the control and tracking system. The position in the arena is recorded at 25 Hz by a camera positioned below a see-through floor. A computer processes the video stream in real time to detect the mouse’s position (centroid) with a custom python script using OpenCV. The software detects when the animal switches between contiguous trapezoids surrounding the towers (see inset in computer screen). Depending on the protocol and the animal’s behavior, specific trapezoid switches trigger the delivery of a single drop of water via one of the four water ports (N, S, E, W) located on the sides of the corresponding tower. Reward delivery at a specific water port of a specific tower is controlled by a Raspberry Pi, which receives commands from the computer and independently activates one of 16 solenoid valves. The labelling of the towers (NE, NW, SE, SW) corresponds to a view from below, which explains why East and West seem reversed on the schematic representation of the arena.

A closed-loop system based on real-time video tracking of the animal positions controls the delivery of drops of water based on the trajectory of the mice (Figure 1C). To ensure that mice obtained a drop of water when moving from one water port to the next (i.e., when turning around a tower corner), we defined four trapezoidal regions (referred to as trapezes) around each tower, each containing a water port. A single drop of water was delivered each time a mouse crossed the boundary between two adjacent trapezes (referred to as a trapeze switch; Figure 1C, see inset on the computer screen). With this basic rule for obtaining rewards and the spatial configuration of the set-up, mice can obtain a series of water drops by circling along the walls of a given tower, going from one water port to the next (i.e., they can *harvest* a tower by performing quarter-turns around it). A critical feature of the task is that when mice begin harvesting a tower, the number of available water drops is not infinite (mice deplete the tower). Therefore, the task discourages mice to only circle around a single tower and requires them to regularly switch towers (explore). Critically, a Python-based control software integrates information about the mice trajectory to further control the water delivery based on: 1) the harvesting direction (e.g., whether the mouse performs quarter-turns in clockwise or counterclockwise direction); 2) which tower can potentially be harvested (e.g., all or a subset of towers deliver water with similar or distinct rules of depletion); 3) ongoing harvesting history (e.g., number of drops obtained or number of quarter-turns performed between ports while harvesting a given tower); and 4) in-between towers history (e.g., which tower can be harvested after leaving a tower, how long after leaving a tower can reward be obtained in the other towers). The combination of these features allows experimenters to study decision-making in a variety of foraging contexts (e.g., different modes of patch depletion and exploration rules) while also engaging processes related to motor skills and spatial cognition.

To illustrate the potential of the TFP, we present two series of experiments conducted on large cohorts of mice. In the first group (Group 1, n=19), we examined whether mice could forage under a fixed procedural constraint: only one turning direction was rewarded. All towers could deliver water, and each time a mouse began harvesting from a tower, we randomly set the maximum number of drops available in that tower for that visit. Once this number was reached, no more drops could be obtained forcing mice to explore. We quantified the mice ability to perform the right action (preference for the quarter-turns in the rewarded direction), their exploration-exploitation tradeoff and the kinematics of their reward-oriented movements. Then we examined how these mice reacted to a sudden reversal of the rewarded direction, both in terms of decision-making (direction preference) and motor control. In the second group (Group 2, n=32), we examined whether mice could forage while the rewarded direction was reversed every day. Finally, we compared the performance of these two groups when submitted to a novel and much uncertain foraging protocol (the rewarded direction and tower is continuously and randomly changing during the session).

### Experiment 1: Stable unidirectional harvesting followed by reversal in Group 1 mice

Mice (n=19, 10 females) were first accustomed to the experimenter during two daily 5-10 minutes handling sessions over 5 days. They were then familiarized with the apparatus during two 15 minutes long sessions (morning and afternoon familiarization sessions, day 1). After the afternoon familiarization session, they were placed under a water-restriction schedule (see methods). Starting on day 2, they performed two daily 12 min-long foraging sessions (one in the morning and one in the afternoon, spaced by at least 4 hours), during which they could obtain drops of water by going from one water port to the next one. Critically, the mice had to adapt to the two following rules. First, only counterclockwise (CCW, from the camera viewpoint) turns around the towers triggered reward delivery at the next water port. Second, each time mice approached a tower, a random number (between 4 and 12, equiprobably) of available drops was randomly assigned for this tower and this visit. Mice could leave a tower at any time, regardless of whether it was fully or partially depleted. After leaving a tower, rewards could only be obtained at the remaining three towers. The rewarded direction was maintained for 6 days (i.e., 12 CCW sessions) after which it was reversed to clockwise (CW) for three days (i.e., 6 CW sessions).

During the familiarization sessions, mice spent more time near the border of the arena and scarcely explored its center (Figure 2AB, left). After a few foraging sessions, mice spent more time turning around the towers (Figure 2AB, center, right). We quantified this trend by computing the difference between the time spent in the trapezes surrounding the towers and the time spent at the border of the arena during the first three minutes of the session (Figure 2C). Overall, mice progressively spent more time around the towers relative to the arena’s border across sessions (Figure 2DE), consistent with increased proficiency in harvesting water.

**Figure 2.**
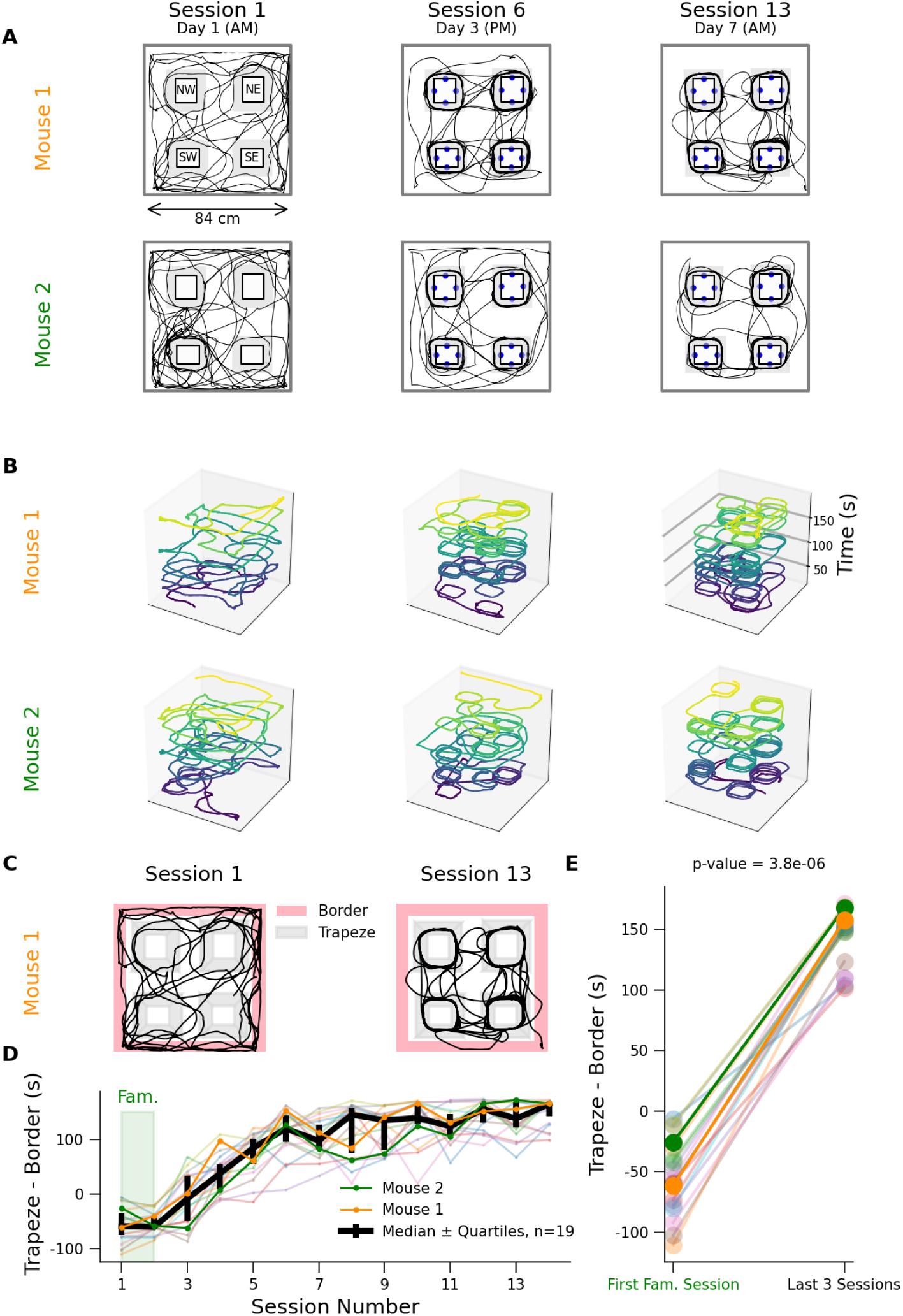
Evolution of group 1 mice trajectories and arena occupancy across sessions. **A**) 2D trajectories of two mice engaged in the Towers Foraging Park (first 3 minutes of sessions 1, 6, and 13; view point of the camera). **B**) 2D trajectories with time extended on z-axis during the first 3 minutes of the same sessions and animals as in A. **C**) Comparison of time spent in the border and trapeze areas for Mouse 1 during 3 minutes of sessions 1 and 13. **D**) Evolution of the trapezes-border time difference across sessions. **E**) Comparison of trapezes-border time difference between the first familiarization session and the last three sessions (n=19).

Video recordings revealed that mice performed discrete runs around and between the towers. We therefore analyzed how these actions (their number and kinematics) evolved across foraging sessions. “Runs” were defined as continuous epochs of movement above a low speed threshold, flanked by immobility periods (Figure 3A; see Methods for details). This definition is agnostic to the type of locomotion (walking, trotting, galloping), although repeated video observations indicated that mice walked. Quarter-turns (QTs) were defined as detected runs starting and ending near two contiguous water ports, shorter than 30 cm and with a duration less than 2 seconds (Figure 3A, red, see Methods for justification of these criteria). Runs Between Towers (RBTs) were defined as runs starting and ending near the water ports from two different towers (Figure 3A, green).

**Figure 3.**
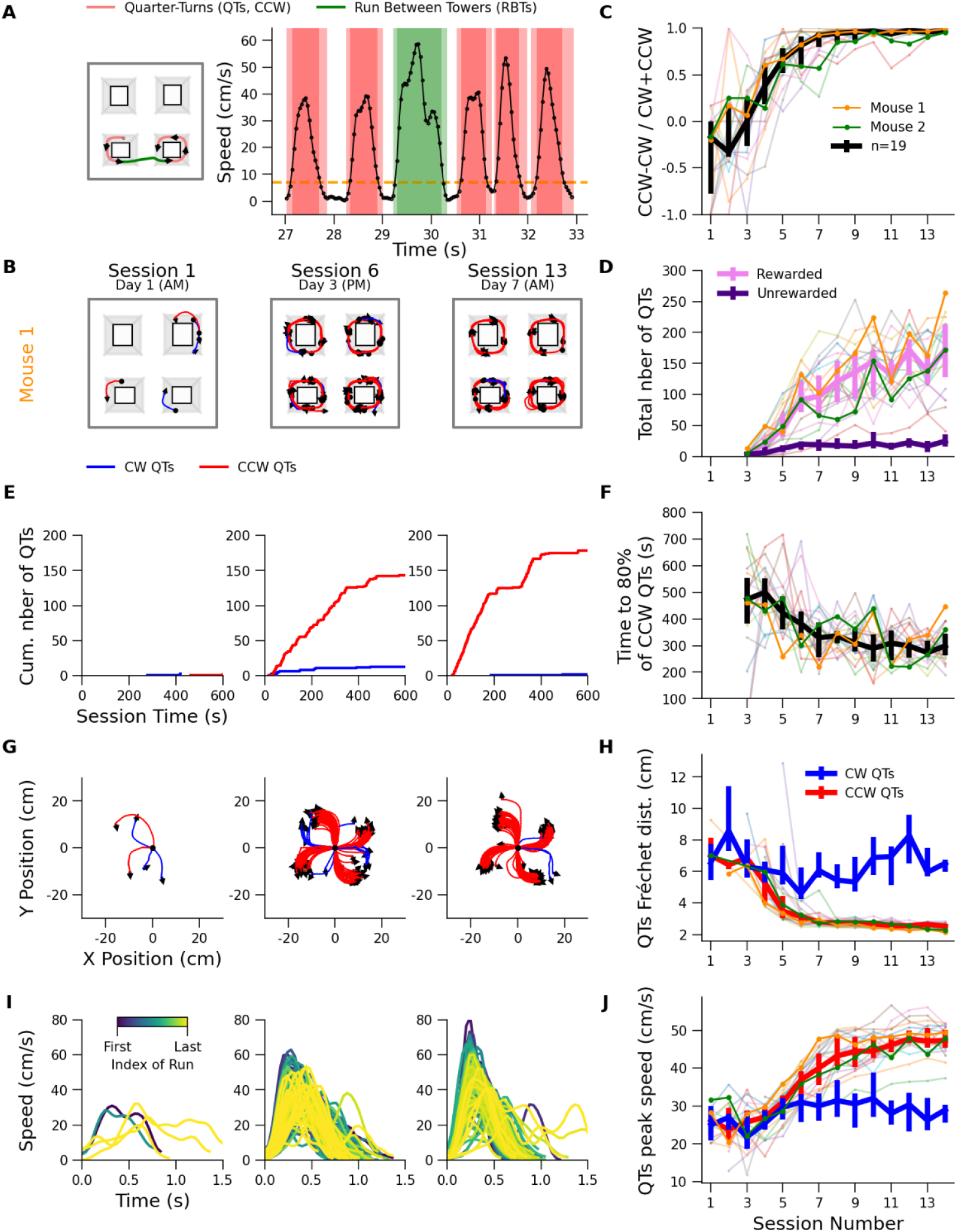
Mice quickly learn to perform quarter-turns in the rewarding direction while optimizing their kinematics. **A)** Short epoch showing the trajectory (left) and speed (right) of a foraging mouse divided into Quarter-Turns (QTs) and Runs Between Towers (RBT). **B)** Detected QTs of Mouse 1 for sessions 1, 6, and 13. **C)** Normalized directional preference of QTs across sessions and all group 1 mice (n=19). **D)** Total number of rewarded (pink) and unrewarded (purple) QTs per session for all the mice. For this group subplot, and all those that follow, thin lines represent individual mice, thick lines represent the group median, and bars extend from the first to the third quartile of the group distribution. **E)** Cumulative number of rewarded and unrewarded QTs during sessions 1, 6 and 13. **F)** Time needed for mice to perform 80% of rewarded QTs across sessions. **G)** Similar to B but QTs share the same origin. **H)** Median Fréchet distance between pairs of CW (blue) and CCW (red) QTs trajectories (rotated and aligned relative to a single tower corner, see Methods) performed during each session. Lower values indicate less variable trajectories. **I)** Speed profiles of QTs during the same sessions as B and E, color-coded according to their rank during the sessions from early (dark blue) to late (yellow). **J)** Evolution of median QTs peak speed across sessions, separately for CW (blue) and CCW (red) QTs.

We first examined how the number, direction, and kinematics of QTs evolved in Experiment 1 (day 1 to day 7; day 1: familiarization sessions; day 2 to day 7: CCW-rewarded sessions). Plotting the detected QTs for sessions 1 (AM, day 1), 6 (PM, day 3), and 13 (AM, day 7) revealed a selective increase in the number of CCW QTs, while CW QTs remained rare (Figure 3B,E,I). To quantify this at the group level, we computed, for each mouse and each session, the normalized difference between the number of CCW and CW QTs. This analysis showed that within just four sessions (across days 2 and 3), all mice developed a strong preference for performing CCW QTs (Figure 3C). If mice keep harvesting a tower after it is depleted, CCW QTs are no longer rewarded. We thus examined how the number of rewarded and unrewarded QTs evolved across sessions. This revealed that mice progressively obtained more rewards without increasing the number of unrewarded QTs (CW QTs or CCW QTs after depletion; Figure 3D). Finally, we examined when QTs occurred during the sessions. The cumulative distribution of CCW and CW QTs performed by the example mouse across the three illustrative sessions showed a growing bias toward CCW QTs, as well as a faster accumulation of CCW QTs at the start of the session (Figure 3E). To quantify this trend at the group level, we computed during each session the time it took for each mouse to perform 80% of their QTs in the rewarded direction. This showed that, with training, mice accumulated rewards at a progressively higher rate (Figure 3F). Altogether these results indicate that mice became progressively more proficient at performing QTs selectively in the rewarded direction without overharvesting the towers.

Aligning QTs to their starting points further illustrated mice’s selective performance of CCW QTs and suggested that trajectories became less variable across sessions (Figure 3G). To quantify this, we realigned and rotated all QTs in a given direction (CCW or CW) from each session relative to a single corner (see Methods). We then computed the Fréchet distance, a measure of similarity between two curves, between all QT pairs in the same direction within each session (see Methods for details and rationale). This analysis showed that CCW QT trajectories became progressively more similar (less variable) across sessions (Figure 3H, red). Noticeably, non-rewarded CW QTs did not become more similar across sessions (Figure 3H, blue). This difference was highly significant and was not due to the smaller number of CW QTs, compared to CCW ones (Figure S1).

Next, we computed the speed profile of the QTs. For the illustrative sessions of the example mouse, this revealed that the speed of QTs increased (Figure 3I). Performing this analysis for CCW and CW QTs across all mice and sessions revealed a progressive increase in the speed of (reward-oriented) CCW QTs, while CW QTs speed remained lower (Figure 3J). This difference was significant and not due to the difference in number of CW and CCW QTs (Figure S2). Finally, for the illustrative mouse, color coding the speed profiles of the CCW QTs according to their order of occurrence within a given session indicated that their peak speed progressively decreased, suggesting that it was influenced by satiety or tiredness (Figure 3I, Morvan, Eloy, and Robbe 2024). Across animals, this effect was observed for a variable proportion of sessions (Figure S3). Altogether, these results indicate that mice quickly learned to execute QTs selectively in the rewarded direction, with trajectories becoming less variable and faster across sessions, and overall they accumulated rewards quicker during each session.

After examining how QTs evolved during learning, we conducted a similar analysis on runs between towers (RBTs). We first plotted the RBTs for a single mouse (same as in Figure 3) during sessions 1, 7, and 14. This mouse performed progressively more RBTs (Figure 4A). Aligning the RBTs at their origin for the illustrative sessions (Figure 4B) and plotting their speed profiles (Figure 4C) suggested that the kinematic parameters of these actions changed across sessions. Indeed, at the group level, we observed that not only did the number of RBTs increase (Figure 4D, left), but they were also performed faster (Figure 4D, center), and their distance decreased (Figure 4D, right). These results indicate that mice performed more RBTs in a faster and more direct manner, most likely contributing to the faster accumulation of rewards during the sessions.

**Figure 4.**
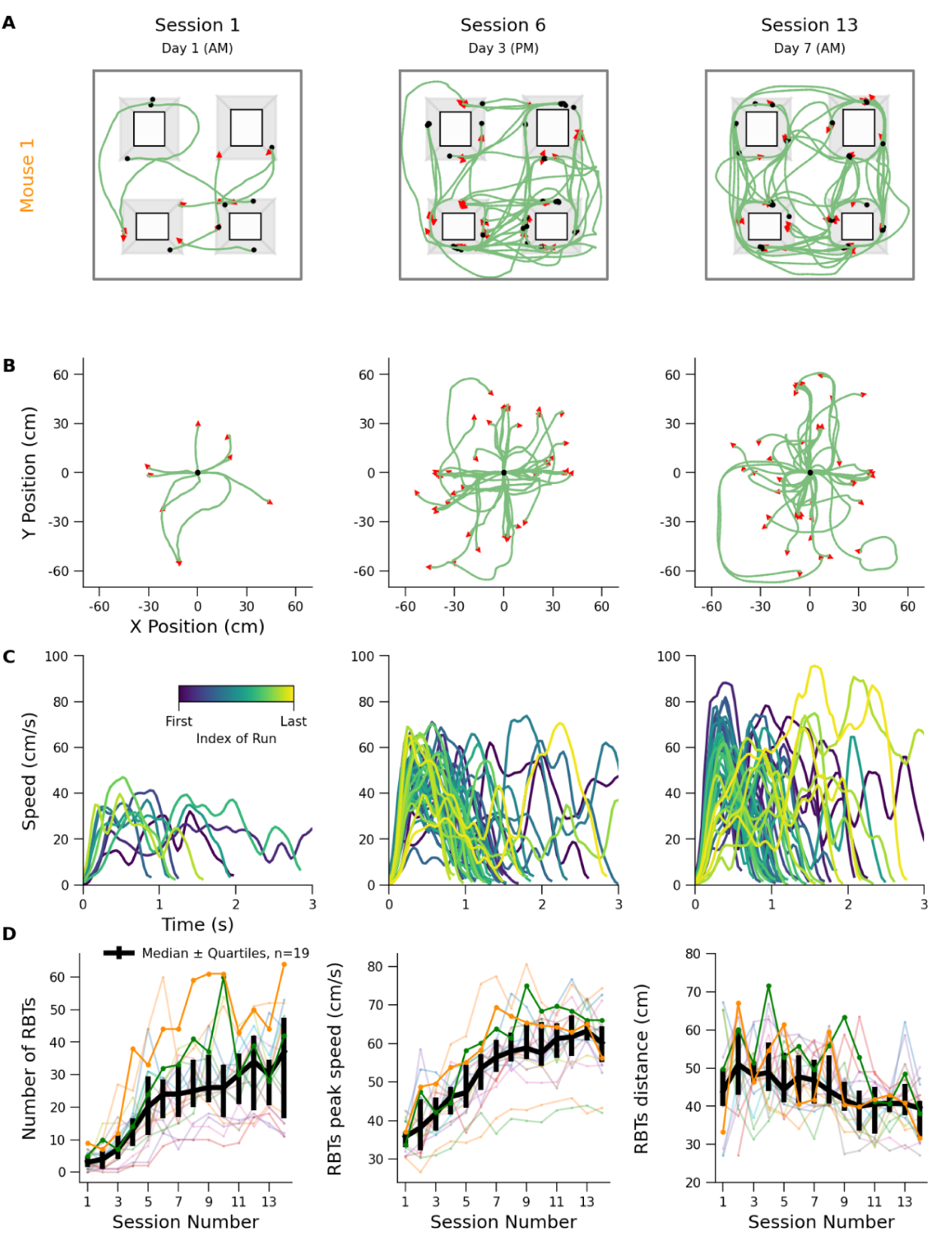
Emergence of fast and short runs between towers. **A)** Detected RBTs of Mouse 1 for sessions 1, 6, and 13. **B)** Same as A but all RBTs share the same origin. **C)** Speed profiles of RBTs for the same sessions as above, color-coded according to their order during the sessions from early (dark blue) to late (yellow). **D)** Across sessions evolution of the number of RBTs (left), their median peak speed (center) and their median length (right), for all the group 1 mice.

The results above are consistent with mice optimizing their behavior (i.e., performing more frequent, faster, and more stereotyped QTs in the CCW direction and switching towers more efficiently) across sessions. For the two illustrative mice, representing the sequence of QTs and their respective location as a raster plot, revealed a large variability in the number of QTs performed at each visit, with many exploitation bouts (consecutive QTs performed around a tower) that were short and for which mice switched tower before performing a non-rewarded QTs (Figure 5A). To quantify how much mice harvested a tower before switching to another one, we computed the number of QTs performed per rewarded visit across mice and sessions.

**Figure 5.**
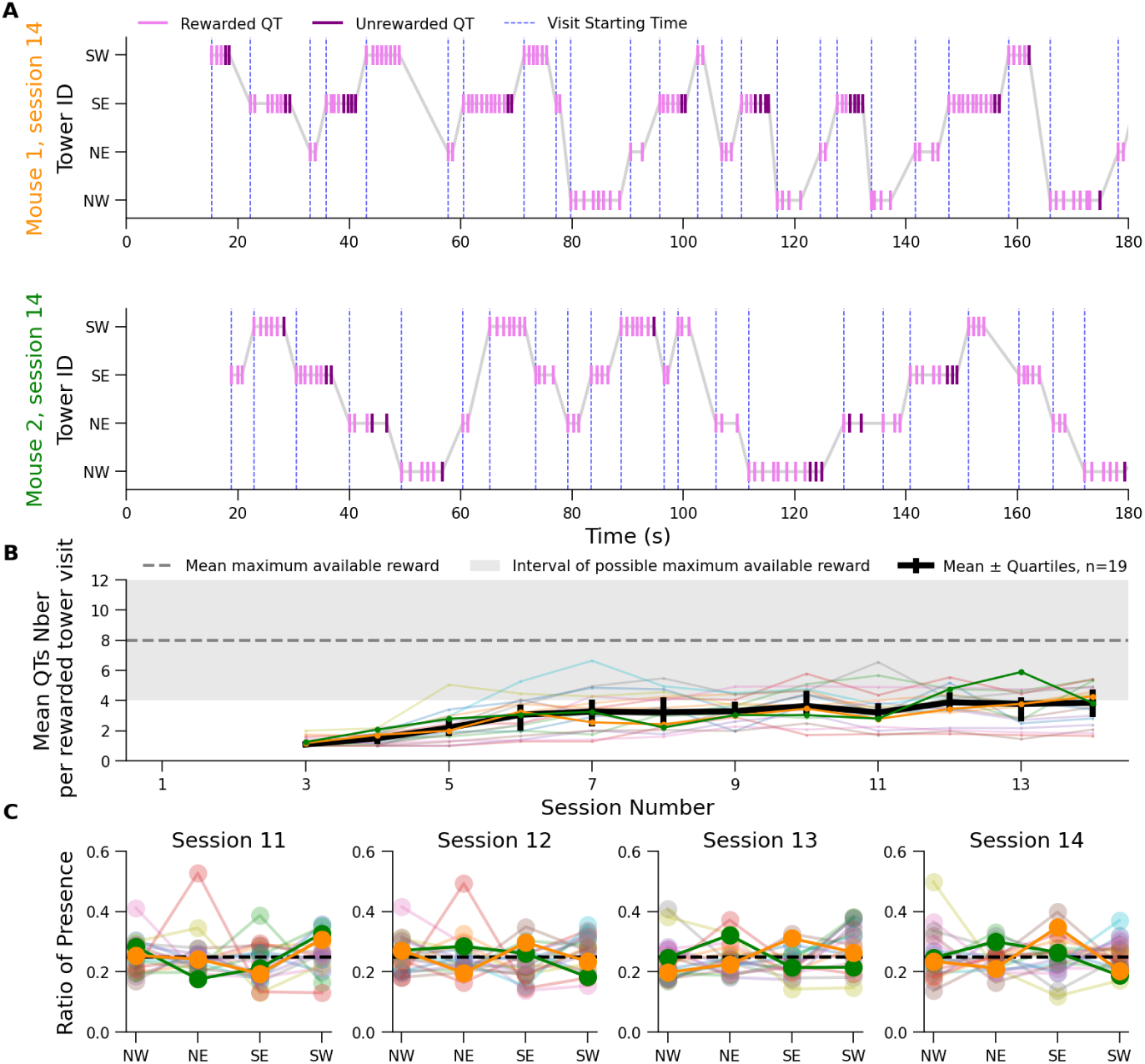
Mice performed short exploitation bouts and homogeneously explored the 4 towers. **A**) Sequence of QTs sorted by tower in which they occur for the two example mice during session 14. **B**) Mean number of QTs per rewarded visit (i.e., visits with at least one rewarded QT) across sessions for all group 1 mice. **C**) Ratio of time spent around each tower for sessions 11 to 14, for all mice.

Although this number increased across sessions, it remained below the theoretical average of available rewards per visit (i.e., 8; Figure 5B). This tendency for short exploitation bouts was paralleled with a relatively homogeneous exploratory drive: mice, on average, spent a similar time around each tower (Figure 5C).

The observation that mice stop harvesting the towers before they are fully depleted (i.e., before performing an unrewarded CCW QT) is consistent with risk aversion and makes sense given the short distance between towers. But it also raises questions over the strength of the procedural learning they acquired. It is possible that the procedural rule to be learned (i.e., making CCW QTs around any tower) is so simple that mice can discover it on the fly at the beginning of each session through rapid trial-and-error. In that case, task proficiency would reflect task familiarization rather than procedural learning per se and should remain largely unaffected if the rewarded direction was suddenly switched from CCW to CW.

To test this possibility, we switched the rewarded direction during the morning session of day 8 (i.e., between sessions 14 and 15; hereafter referred to as the directional switch) and examined how mice adapted their behavior. We observed a transient drop in the number of rewarded QTs during this first session post directional switch (Figure 6A, pink). In this session, mice continued to perform nearly as many CCW QTs as in the previous session, despite these actions no longer being rewarded (Figure 6A, violet). This persistence is consistent with the acquisition of procedural memory. Nonetheless, mice did begin to perform CW QTs in this first session post directional switch, albeit at much lower rate than CCW QTs (Figure 6AB). Strikingly, by the following afternoon session, nearly all mice had already reached a performance level comparable to that of previous days. We also examined how the rule reversal affected the time required for mice to accumulate rewards. As expected, mice took much longer to obtain 80% of their rewards during the first session after the direction reversal (Figure 6C), but this time rapidly decreased in subsequent sessions. This post-reversal improvement occurred faster than during initial learning, suggesting a form of meta-learning. Overall these results indicate that mice acquired a form of procedural memory during the first 6 days of training but can update it quickly in response to a change of rule.

**Figure 6.**
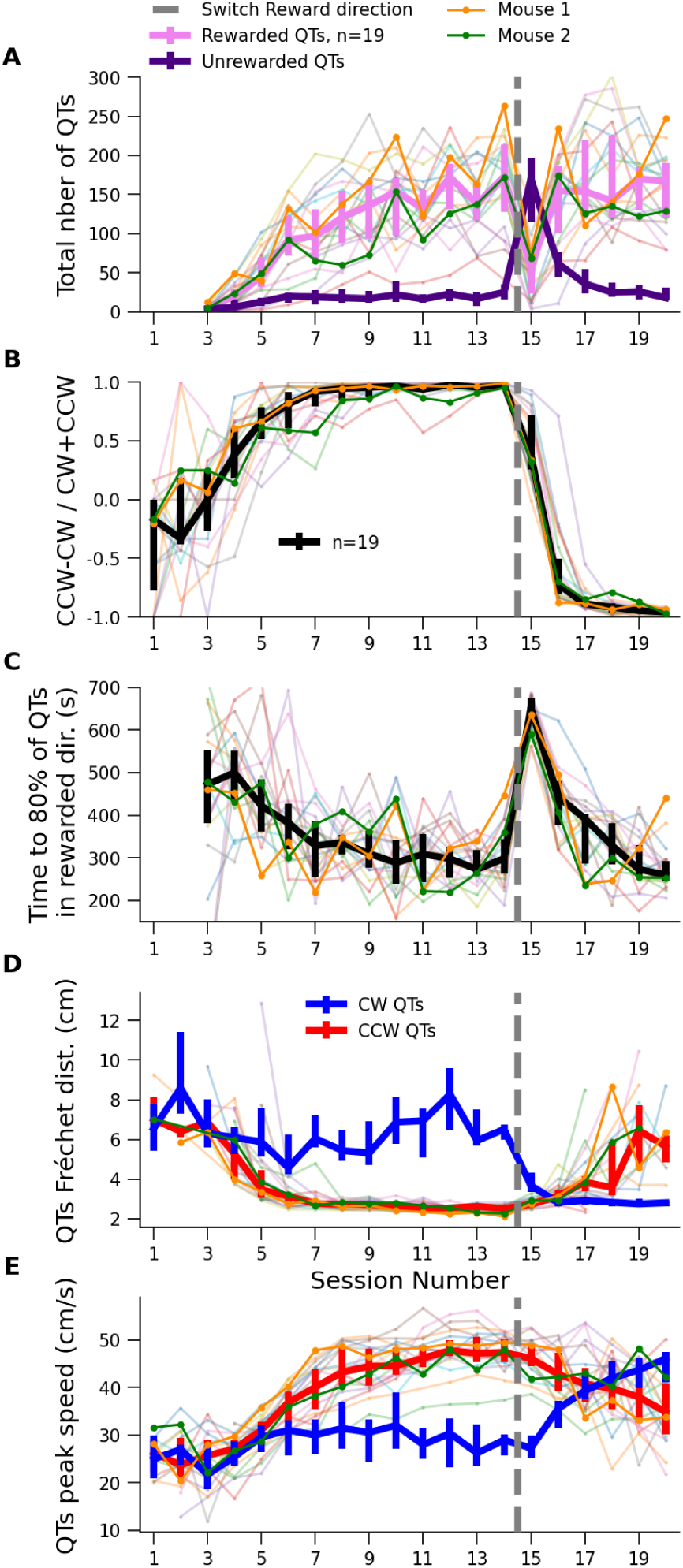
Foraging adaptation following abrupt change of the rewarded direction. **A)** Evolution in the number of rewarded (pink) or unrewarded (purple) QTs throughout the sessions, before and after change of rewarded direction (dash grey line) for group 1 mice. **B)** Similar to A for QT’s normalized directional preference. **C)** Similar to A for the time necessary to gather 80% of rewards. **D)** Similar to A for QTs trajectory variability measured by the median Fréchet distance across all pairs of CW (or CCW) QTs. **E)** Similar to A for QTs speed.

Next, we examined the impact of the rule reversal on the kinematics of CCW (previously rewarded, now unrewarded) and CW (previously unrewarded, now rewarded) QTs. During the first post-reversal session, mice performed only a few CW QTs (Figure 6A,B) but their variability was already much lower than in the pre-reversal sessions, quickly reaching its minimum in the following sessions (Figure 6D, blue). Such a rapid change in movement variability after rule reversal closely mirrors the change in QT direction preference (compare black line in Figure 6B with blue line in Figure 6D). Interestingly, the variability of CCW QTs progressively increased after the rule reversal (Figure 6C, red). Together, these findings indicate that QTs variability is highly sensitive to recent reward history. On the other hand, the speed of CW QTs did not increase during the first post directional-switch session (Figure 6E, blue) and mice continued performing unrewarded CCW QTs with the same fast speeds as in the previous session (Figure 6E, red). In addition, during the following post directional-switch sessions (sessions 16–20), the speed of CW QTs slowly increased even though directional choice selectivity had already plateaued around 85% by session 17 (Figure 6E, blue). During these same sessions, the speed of CCW QTs slowly declined (Figure 6E, red), suggesting that movement speed reflects the history of reward outcomes over several sessions. Overall, these results showed that the mice’s ability to flexibly update their preferred harvest direction (from CCW to CW) was accompanied by unexpectedly decoupled changes in the variability and speed of the QT trajectories, with the latter changing much more slowly than the former.

### Experiment 2: Daily reversal of harvesting direction in Group 2 mice

In a second group of mice, we further examined their ability to regularly update their procedural preference (performing CW or CCW QTs) when the rewarded direction was switched every day. Similarly to the previous group, these mice were first subjected to two familiarization sessions (day 1) before entering the water-restricted protocol. On day 2, during both morning and afternoon sessions, mice could obtain rewards by performing QTs in both directions. Starting on day 3, the rewarded direction was fixed each day and changed every morning (see methods for details). Across sessions, this group of mice displayed a similar pattern of change in behavior than the previous one: mice started by exploring primarily the outer border of the arena but quickly spent more time running around the towers (Figure S4). Examining the QTs performed by a single mouse across 5 illustrative sessions (session 1 on day 1, sessions 9 to 12 on days 5 and 6) showed that, by day 5, the mouse performed many fast QTs (Figure 7A-E). However, during the illustrative morning sessions (sessions 9 and 11), this animal performed QTs in both rewarded and unrewarded directions, while in the afternoon sessions (sessions 10 and 12), it mainly performed QTs in the correct rewarded direction (CCW and CW for session 10 and 12, respectively, Figure 7B-D). Examining the cumulative number of CCW and CW QTs from session 9 to 12 revealed that in the morning sessions, the mouse started by performing QTs in the direction rewarded the previous day and progressively performed less of these non-rewarded turns to switch to rewarded direction. In contrast, during the afternoon sessions, mice immediately performed QTs in the rewarded direction and very few QTs in the unrewarded direction (Figure 7C).

**Figure 7.**
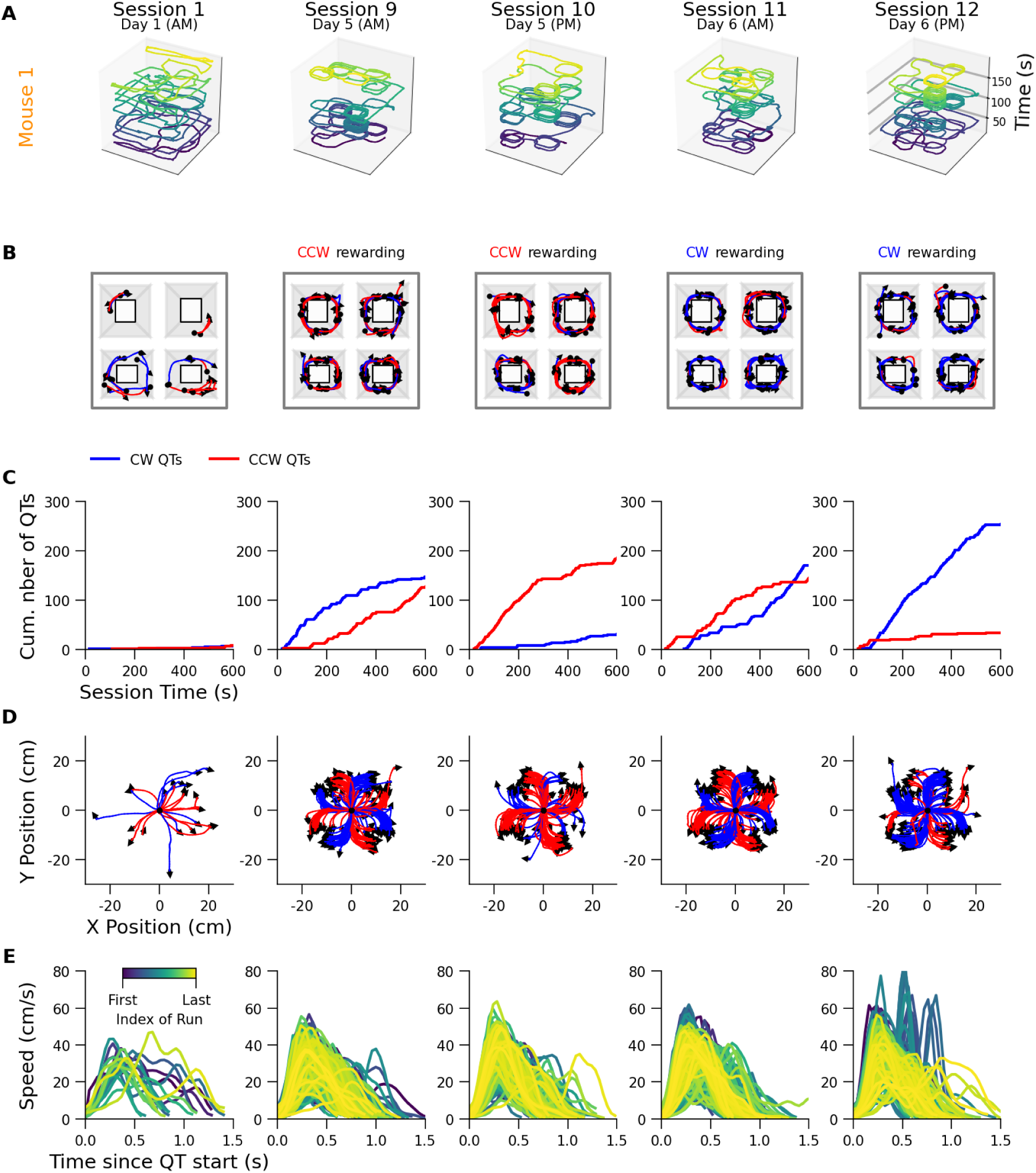
Learning and adaptation of a single mouse when the directional rule is reversed every morning. **A**) 2D trajectories with time extended on the z-axis for 5 sessions (first 3 minutes; session 1 on day 1 and 9 to 12 on days 5 and 6) for a single individual from group 2 mice. **B)** Detected QTs for the same mouse and sessions as A. **C)** Cumulated number of CW and CCW QTs during the same sessions. **D)** Same as B but all QTs share the same origin. **E)** Speed profiles of QTs during the same sessions, color-coded according to their rank during the sessions, from early (dark blue) to late (yellow).

Such a behavior was observed for an entire population of mice (n=32 mice, 12 females). Indeed, plotting the normalized preference for CCW and CW QTs revealed that mice systematically adapted their QTs direction choices (CW or CCW) according to the daily rewarded direction, although the bias was systematically less pronounced in the morning (Figure 8A). The same result was observed by plotting the number of CCW and CW QTs across sessions (Figure 8B). It also confirmed that mice performed less (more) rewarded (unrewarded) QTs in the morning sessions compared to the afternoon sessions. However, this difference diminished with training. By days 9 and 10, mice performed, on average, the same number of rewarded CW (or CCW) QTs in the morning and afternoon sessions (compare blue traces between sessions 17 and 18, and red traces between 19 and 20; Figure 8B). Thus, mice display meta-learning when repetitively challenged with reward directions reversals. In addition, mice were progressively more efficient to perform CW and CCW QTs across sessions although, as expected, they took longer to accumulate rewards in the morning compared to afternoon sessions (e.g., on a CCW rewarded day, mice to obtain rewards by performing CCW QTs in the morning than in the afternoon session, Figure 8C). Still, mice were progressively more efficient in the morning to perform QTs in the rewarded direction, consistent with meta-learning.

**Figure 8.**
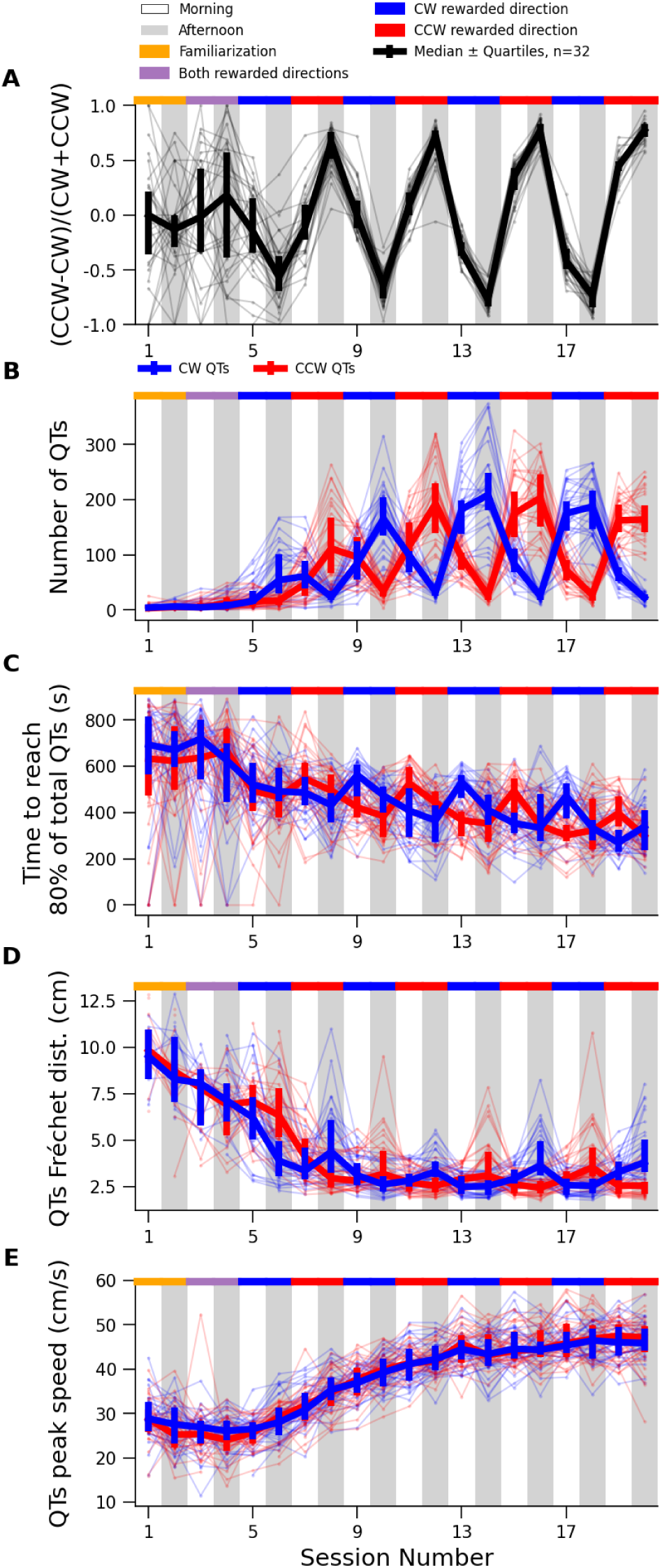
Decision and kinematic adaptation to daily fluctuation in rewarded direction. **A**) Normalised directional preference of QTs across sessions for group 2 mice (n = 32). **B**) Number of CW and CCW QTs across sessions and mice. **C**) Time necessary for mice to perform 80% of CW (blue) and CCW (red) QTs across sessions. **D**) Median Fréchet distance of CW (blue) and CCW (red) QTs trajectories per session, across sessions and mice. Lower values indicate less variable trajectories. **E**) Median peak speed of CW (blue) and CCW (red) QTs trajectories per session and animals, across sessions.

Next, we examined the kinematics of QTs across sessions in this group of mice. The reduction in QTs trajectory variability developed quickly and varied with the rewarded direction (Figure 8D). Specifically, the Fréchet distance between CCW QTs decreased when CCW turns were rewarded and increased when CW turns were rewarded. A symmetric modulation was observed for CW QTs. In contrast, even if mice gradually performed both CCW and CW QTs faster (Figure 8E), this change was slower than the decrease in trajectory variability, confirming that the speed of QTs integrates the history of reward over several sessions. In line with this interpretation, the speed of both types of QTs (CW or CCW) did not fluctuate across days (i.e., according to the rewarded direction).

Finally we examined changes in RBTs across sessions for this second group of mice. Similarly to the first group of mice, a progressive increase in the number and speed of RBT, and a reduction in their distance, were observed (Figure S5). In addition, we observed a tendency for mice to perform more RBT in the morning than in the afternoon session, which suggests that, when they harvest the tower in the wrong direction, mice primarily change towers rather than switch direction. Overall, in this second group of animals, we observed performance improvements (preference for QTs in the rewarded direction, QTs kinematics’s overall improvement and modulation by reward history, RBTs number and kinematics) that were highly consistent with those seen in Group 1, despite daily reversals of the rewarded direction. Critically, Group 2 mice adapted increasingly quickly to these reversals, consistent with the emergence of meta-learning.

### Experiment 3: Adaptation of Group 1 and Group 2 mice to random foraging rules

Finally, we examined how the two groups of mice reacted when, after adapting to the directional switch (either only once for Group 1, or daily over several days for Group 2), they were challenged with a highly uncertain foraging condition in which both the rewarded direction (CW or CCW) and the identity of the next rewarded tower (NW, NE, SE, or SW) were randomly reassigned each time a mouse left a depleted tower (hereafter referred to as uncertain protocol, see methods). The number of rewarded QTs dropped when Group 1 mice were exposed to the uncertain protocol (Figure 9A, left, pink). An opposite modulation was observed for unrewarded QTs, which abruptly increased during the first sessions of the uncertain foraging protocol (Figure 9A, left, purple). This inefficiency was not due to the mice persevering in the previously rewarded direction. Indeed, the CW bias inherited from previous foraging sessions was reduced during the first session post-protocol change and totally nonexistent after three sessions in this new protocol, suggesting that mice from Group 1 struggled to quickly find the rewarded tower and direction rather than flexibly performing CW or CCW QTs (Figure 9B, left). In striking contrast, the second group of mice’s number of rewarded QTs hardly dropped in the first session under the uncertain rule (Figure 9A, right, pink). Thus, even if their number of unrewarded QTs logically increased in a protocol in which mice cannot guess which tower and direction will be rewarded when they leave a depleted patch (Figure 9A, right, purple), they ultimately managed to perform a similar number of rewarded and unrewarded QTs. Moreover, Group 2 mice immediately reduced their CW directional bias from the previous session, performing as many CW as CCW turns as early as the first session under the uncertain protocol (Figure 9B, right). The better performance of Group 2 mice compared to Group 1 (limited decrease in rewarded QTs), and their quicker adaptation to the lack of directional bias in the uncertain foraging protocol could be observed in individual mice (Figures S6 and S7) and was statistically significant (Figure S8). Mice from Group 1 therefore obtained fewer rewards than mice from Group 2. In both groups, the speed and variability were similar for CW and CCW QTs, as expected as in this uncertain condition both of them have the same probability to yield reward (Figure 9CD), confirming again the dependency of these two features to reward history.

**Figure 9.**
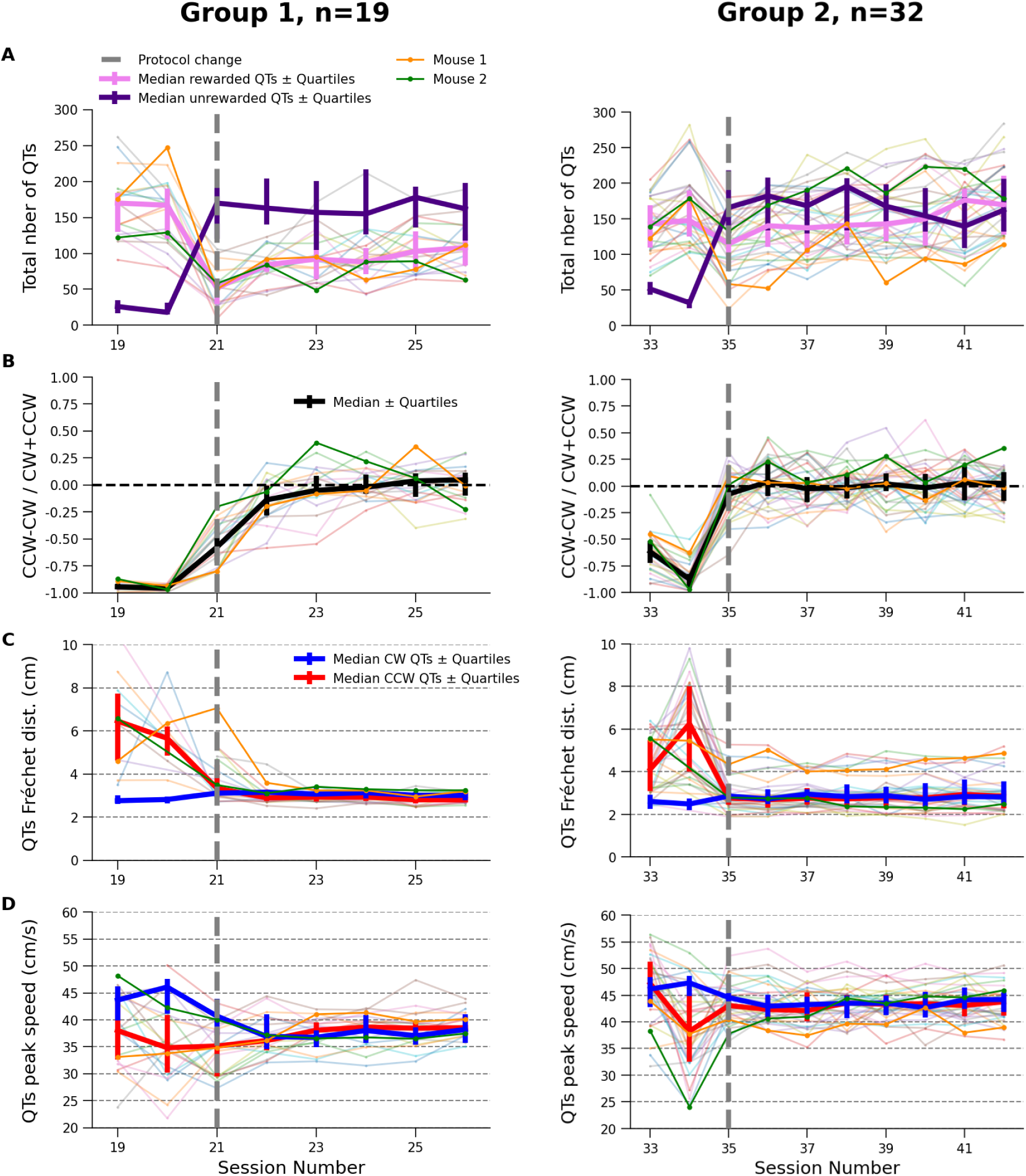
Impact of prior exposure to directional switches on adaptation to uncertain foraging. **A)** Number of rewarded (pink) and unrewarded (dark blue) QTs across sessions for group 1 (left) and group 2 (right) mice. Dashed gray line indicate first session under uncertain protocol. **B)** Evolution of directional bias (CW vs. CCW QTs) across sessions. **C)** Trajectory stereotypy of CW and CCW QTs, measured by the median Fréchet distance across sessions for group 1 and group 2 mice **D)** Median Peak speed of CW and CCW QTs across sessions for group 1 (left) and group 2 (right) mice

In the uncertain protocol, the time required to find water after leaving a depleted tower necessarily increases, as the rewarded tower and direction are randomly reset. According to the marginal value theorem (Charnov 1976), when foragers seeking to maximize their reward rate face longer exploration times, they should increase the time spent harvesting patches. While our uncertain protocol does not fully conform to the assumptions of the marginal value theorem, it still suggests that mice should prolong their harvesting activity under these conditions. Indeed, we found that both groups immediately increased the number of QTs per rewarded tower visit (Figure S9). Interestingly, mice from Group 1, when subjected to the uncertain foraging protocol, displayed a trend toward more QTs during unrewarded tower visits, compared to Group 2 mice (Figure S10). Finally, that Group 2 mice displayed greater capacity to adapt to the continuously changing rewarded location and direction, compared to Group 1, was also confirmed by an increased number of RBTs in the uncertain condition for Group 2 (compare Figure S11 et Figure S12). Altogether, these results indicate that, compared to Group 1 mice, Group 2 mice spend more time exploring and less time perseverating on unrewarded harvesting attempts.

## Discussion

In this study, we present experiments conducted in a newly developed behavioral framework that recreates the dilemmas animals face during patch foraging. The results show that mice rapidly developed adaptive foraging strategies. They harvested the towers in the rewarded direction during short bouts of exploitation and, across sessions, the speed and variability of their reward-oriented movements (but not of non-rewarded ones) increased and decreased, respectively. Our experimental paradigm not only allows us to study learning, decision-making, and motor control but also reveals the animals’ capacity to adapt to an unexpected change in procedural rule (harvesting the towers in a given direction). We found that mice quickly, though not instantaneously, switched their turning direction and displayed an intriguing decoupling of the two main kinematic features of their reward-oriented movements: they modified their trajectories faster than their speed. In addition, mice subjected to several reversal of the rewarded direction became faster at adapting, revealing a new form of meta-learning. Critically, we found that these mice performed better when subjected to a more challenging condition, providing a unique functional demonstration of this adaptive mechanism. Below, we first discuss why our task can be considered naturalistic and compare it to previous attempts to model the exploration-exploitation trade-off. We then elaborate on the two main mechanistic findings of the study: the decoupling of speed and trajectory, and meta-learning, to highlight its potential to understand how animals flexibly adapt to unexpected changes in their environment.

### The Towers Foraging Park, a naturalistic behavioral test ?

Here we present the Towers Foraging Park, an experimental framework designed to study learning, reward-guided decisions, and movements in freely moving mice. This experimental development should be seen in the context of a growing interest in studying behavior under more naturalistic conditions (El Hady et al. 2019; Dennis et al. 2021; Rosenberg et al. 2021; Cisek and Green 2024; Oesch et al. 2024; Ulanovsky 2025). Defining what qualifies as naturalistic is a complex issue (Chrzanowska et al. 2025). We take this challenge as a starting point to outline key features that, in our view, make our task ethologically sound, despite the inevitable constraints of studying behavior in a laboratory. First, mice were allowed to move freely in a large arena (relative to their body size) and the task was not divided into trials and intertrial periods. Second, the task requires mice to forage along walls in complete darkness, which aligns with their nocturnal nature and leverages on their spontaneous tendency to perform thigmotaxis (Leppänen et al. 2006). Third, the task did not require mice to associate arbitrary instrumental responses (e.g., lever presses) or neutral stimuli (a sound, a light) with the remote delivery of rewards. Instead, mice directly checked for the presence of water drops at the tips of salient water ports located on the sides of the towers. Critically, our custom software continuously tracked the position and the history of reward-oriented actions to implement reward delivery rules according to the behavior of the mice and with built-in variability. This variability constitutes the fourth feature that makes our task ethological: after a small, randomly determined number of rewards, the towers became transiently depleted. This mimics two key features of natural environments, namely, that repeated actions do not always yield the same outcomes, as resources tend to deplete or fluctuate over time, and that mice have several options to obtain rewards.

Clearly, every scientific attempt to understand the living comes with inherent limitations. There will always be a gap between “real-life” experience and the simplifications (or abstractions) required to study brain functions scientifically (Chirimuuta 2024). Our new task is no exception to this limitation. First, mice were motivated to forage through a controlled water restriction schedule. This introduces an artificial element and some stress, although it is worth noting that hunger and thirst are not unnatural states for wild animals. Unlike humans, who store food and water at home, most animals, including rodents, must seek out resources on a regular basis. Thus, although thirst was experimentally induced, the resulting drive to forage reflects part of their innate behavioral repertoire. Second, and related to the first point, mice were placed in the arena by an experimenter, who also returned them to their home cage when the session ended (session duration was fixed at 12 minutes). In the future, we aim to provide mice with on-demand access to the foraging area rather than having experimenters handle the animals at regular times of the day (Torquet et al. 2018). This kind of approach would open the door to a wide range of questions, such as whether the urgency to drink during a limited number of short sessions, and the potential physiological stresses induced by these protocols, affect mice’s decision-making and movements.

As pointed out by Chrzanowska et al. (2025), what qualifies as a “naturalistic” behavioral task is to some extent subjective, and it would be problematic if this term was used as a *petitio principii* or a virtue signal to implicitly dismiss other approaches, especially some that have already yielded key insights into neural underpinning of behavior. Still, given the mounting societal pressure and push against animal experimentation, neuroscience cannot avoid reflecting on how it approaches animal behavior. Indeed, several potential issues can arise when studying brain function during non-naturalistic tasks. The first is that overly constrained experimental conditions may promote behaviors outside the animal’s natural repertoire, leading to the investigation of non-natural brain patterns (Krakauer et al. 2017). The second is that such conditions can create distress, an obvious major concern, both ethically and scientifically. The third is the risk that, in unnatural settings, animals may not comply with the anthropomorphic expectations of the experimenter and instead solve the task using idiosyncratic heuristics, or more generally by engaging brain functions other than those originally targeted (Robbe 2023). Our own experience in training mice and rats on trial-based tasks has led us to identify two key parameters that could serve as markers of non-naturalistic design: long training protocols involving multiple progressive phases, and the frequent observation of uninstructed stereotyped movements (rituals) that appear necessary for task completion and that animal have difficulties to unlearn despite change in reward outcome (Rueda-Orozco and Robbe 2015; Sales-Carbonell et al. 2018; Safaie et al. 2020). Several aspects of our data provide empirical grounding for describing the TFP as naturalistic. First, after spontaneously exploring the entire arena with a clear preference for the outer borders, mice rapidly developed a foraging strategy (circling around the appropriate tower), typically within just three to four 12-minutes long sessions (Rosenberg et al. 2021). Second, the kinematics of their reward-oriented movements showed expected signs of utility-based modulation, both across and within sessions. Third, mice maintained a high level of exploration of the different towers interspersed with brief exploitation bouts, which matched the task’s geometry. They did not develop prolonged exploitation nor overly stereotyped sequences of explorations of specific towers. Fourth, despite clear procedural learning, mice quickly adjusted their choices and movements when the task rules changed. Fifth, mice exposed to regular rule changes learned to become more flexible and adapted quickly to drastic changes in foraging uncertainty. Finally, although there was clear individual variability in the above-described behavioral features, the trends we observed were observed across two large groups of 19 and 32 animals, trained in distinct protocols and were robust across sessions (i.e., mice did not suddenly stop working or give up, which is often the case in head-fixed paradigm).

### Comparison with multi-armed bandit tasks in operant boxes

Our task is inspired by patch foraging, a behavior shared by most animals that requires solving the so-called exploration-exploitation dilemma (Stephens and Krebs 1986; Addicott et al. 2017; Barack 2024). In rodents, the exploration-exploitation dilemma is most often studied using multi-armed bandit tasks, in which animals must choose between several similar actions (typically choosing to press on one lever out of 2 or 3) to obtain a reward (typically delivered in a magazine placed on the wall opposite to the levers). Each action is associated with a distinct probability of reward delivery, and these probabilities fluctuate in unsignaled blocks during the session. Exploitation and exploration are typically inferred from the ratio of responses to the high- and low-value action and from the latency to switch lever after reward contingencies are reversed. These tasks have provided major insights into the neural mechanisms that support flexible decision-making in uncertain environments (e.g., Tai et al. 2012; Hamid et al. 2016; Cinotti et al. 2019; Mohebi et al. 2019). Nonetheless, they may not fully capture the richness of the exploration-exploitation dilemma. First, bandit tasks for rodents are usually conducted in small operant boxes, whereas exploration and exploitation are inherently spatial behaviors, especially for animals such as mice and rats. This is important because time and effort are important determinants of economic decision making (Carland et al. 2019; Shadmehr et al. 2019). Second, in the bandit setup, the available options (typically pressing a lever or poking its nose into a hole) are nearly identical in terms of their spatial and motor requirements. This makes it difficult to dissociate exploration from exploitation based on overt behavior alone. For example, pressing the low-probability lever could reflect either an exploratory action (testing whether its value has changed) or an exploitative one (if the animal expects it to yield a reward). Still, pressing the low-reward lever is interpreted as exploration. In our experiments, when the rewarded direction was fixed, mice rarely performed QTs in the non-rewarded direction, and when they did, their movements were slower and more variable than in the rewarded direction. Yet mice were not overly exploitative, as they generally underharvested the patch. These observations highlight the analytical granularity of our experimental design, independent of theoretical models. Third, these tasks typically involve large differences in reward probability between the two levers at any given time. In contrast, real-world foraging dilemmas often involve deciding when to leave a still-rewarding source, not only choosing between a good and a bad one. In the bandit setting, when reward probabilities are swapped, the animal has no choice but to switch, making it unclear whether this reflects genuine exploration or merely the abandonment of an inferior option (Wang et al. 2023). Fourth, in operant versions of bandit tasks, the location of reward delivery is physically separated from the location of the choice (e.g., lever vs. magazine). This separation limits direct comparison to natural foraging problems, such as patch depletion, where exploitation and reward consumption are spatially and temporally coupled. To structure trials and encourage choice behavior, arbitrary cues such as lever retraction, or trial initiation sensory cues are often introduced. This requires conditioning the animals during lengthy pre-training phases which may bias behavior toward exploitation and reduce its natural flexibility. Finally, multi-armed bandit tasks rarely track for the kinematics of reward-oriented movements, they focus almost exclusively on the timing of lever-press and food magazine entry, ignoring the kinematics of the actions themselves even though they convey important information about decision-making (Shadmehr et al. 2019), including during foraging (Yoon et al. 2018).

In contrast, in our setup, mice rapidly engaged in foraging behavior without any pretraining, aside from two short familiarization sessions on day one. The foraging options (towers) are spatially segregated, making both exploration and exploitation kinematically tractable on a 2D plane (e.g., QTs vs. RBTs). Notably, we observed that mice often ceased exploiting a rewarding tower even when rewards were still available, likely an adaptive strategy due to the proximity of neighboring towers. We also found that exploitation behavior was modulated by the duration of prior exploration: under the high-uncertainty protocol, all mice exhibited more prolonged exploitation once they located a rewarding tower/direction (Figure S9). This suggests that their behavior is sensitive to both uncertainty and time, in line with theoretical prediction (Kilpatrick et al. 2021b). Finally, our task can provide a bridge with human and non-human primate patch-foraging paradigms (Hayden et al. 2011; Barack et al. 2024). Indeed, foraging behavior has recently emerged as a valuable model for studying the exploration-exploitation tradeoff and its relationship to mental health (Jami et al. 2025).

### Rule reversal reveals separate effects of reward outcome history on movement speed and variability

The TFP provides an interesting playground to study motor learning and investigate how reward influences motor control. Indeed mice not only learned which action to perform to obtain water (CW or CCW QTs) but also displayed pronounced modulation of how they execute these actions. Thus, our study allows us to probe both action selection and execution, two fundamental components of motor skills (Chen et al. 2018; Krakauer et al. 2019). As mice became more familiar with the task structure and expected to find water around the towers, we observed improvements in both action selection (e.g., animals from Group 1 performed more CCW than CW rewarded trajectories) and movement execution. Specifically, QTs were progressively executed faster and their trajectories displayed less trial-by-trial variability (Figure 3). This phenomenon is well documented for visual saccades or reaching movements in both humans and non-human primates (e.g., Summerside, Shadmehr, and Ahmed 2018; Takikawa et al. 2002). In rodents, reduced variability and increased speed are typically interpreted as signs of automatization of reward-oriented movements or action sequence (Jin and Costa 2010; Rueda-Orozco and Robbe 2015; Lemke et al. 2019; Jurado-Parras et al. 2020; Mizes et al. 2022), reflecting a relatively unified motor learning process. Surprisingly, when mice from Group 1 performed their first session following rule reversal (i.e., switching from six days of training with CCW QTs rewarded to CW QTs being rewarded), we observed a clear increase in the variability of CCW QTs, while their speed remained unchanged (i.e., fast). Analysis of CCW QTs across sessions confirmed that variability increased more rapidly than speed decreased. A similar dissociation was found for CW turns: before the rule reversal, these unrewarded actions remained both slow and highly variable. Yet as early as the first session after reversal, their variability decreased while speed remained low. Over subsequent sessions, variability quickly reached a low plateau, whereas the speed of CW QTs increased more gradually. This pattern suggests that reward-related movement variability is highly sensitive to recent reward history (Dhawale et al. 2019), while speed integrates reward outcomes over a longer timescale spanning several sessions. The differential influence of reward history on QT speed and variability proved robust, as it was also evident in Group 2 mice. For these animals, the rewarded direction was swapped each morning, and the variability of CCW and CW QTs oscillated in sync with this alternation (Figure 6C). In contrast, speed showed no such oscillations and instead rose steadily, consistent with the idea that it reflects reward integration across multiple sessions (Figure 6D).

The above decoupling between the speed and variability of reward-oriented movements is interesting for several reasons. First, if reduced QT variability simply reflected the consolidation of a motor skill, it is unclear why movement variability would increase so quickly once the action became unrewarded. In the rodent literature, the stereotypy of movements is often associated with habitual or skilled action (i.e., robust to change in reward contingencies). Here, the reported high sensitivity of QT variability to reward history echoes works in humans (e.g., Pekny et al. 2015; Manohar et al. 2019) and rats (Dhawale et al. 2019), and is more congruent with an exploration-exploitation tradeoff at the movement level, enabling adaptive behavior when reward outcome fluctuates (Dhawale et al. 2017). Second, reward-dependent movement invigoration is a well-known adaptive process allowing humans and other animals to reduce temporal discounting of rewards (Shadmehr et al. 2019). But this process raises a problem for the nervous system. Indeed stronger motor commands are more susceptible to noise, potentially reducing accuracy, which suggests the need for error-reducing mechanisms. It has been proposed, for example, that reward may compensate not only for the energetic cost necessary to move quickly but also for the cost of maintaining precision, acting therefore on two separate aspects of motor control (Manohar et al. 2015). The reduction of motor noise has been proposed to involve multiple mechanisms including increased limb stiffness (Codol et al. 2020). By establishing a model in which locomotion speed and variability can decouple in response to sudden change in reward history, our study opens the way to investigating the neural mechanisms underlying this dissociation (e.g., distinct brain circuits under the influence of dopamine). Here, for the sake of simplicity, all our analyses were based on tracking the center of mass of the mice. In the future, a better quantification of the entire body position will be incorporated (Mathis et al. 2018; Pereira et al. 2022). Indeed while biomechanical costs have been shown to influence motor decisions in humans (Cos et al. 2011; Canaveral et al. 2024), the extent to which similar modulation contributes to foraging behavior is, to the best of our knowledge, largely unknown, despite obvious relevance in the context of a freely-moving foraging task (Cisek and Pastor-Bernier 2014).

### Repetitive reversion of harvesting direction reveals functional meta-learning

An interesting observation from our experiments is that Group 2 mice, which were exposed to daily reversals of the rewarding direction, became progressively faster at updating their choice in the morning sessions (Figure 8). In these sessions, Group 2 mice began by harvesting in the direction rewarded on the previous day (see Figure 6C), showing that the animals relied on their past experience to forage. This persistence of mice in performing QTs in a non-rewarded direction suggests that they had formed a “belief” that this direction should yield reward, and required repeated contradictory outcomes to revise their policy. This strategy is adaptive, since the absence of reward could result from transient noise, and it would be suboptimal to abandon a previously successful strategy after a single failure. However, excessive rigidity would be detrimental. Strikingly, across sessions, Group 2 mice performed fewer unrewarded QTs and more rewarded QTs during the morning sessions (Figure 8AB, see also the gap between red and blue curves in session 11 cs session 9 in Figure 7C). This translated into progressively faster reward accumulation (Figure 8C), indicating that the mice adapted more quickly to the daily change in reward direction. If this improvement reflects a form of meta-learning, it should provide an advantage in different task versions, compared to Group 1 mice trained under more stable contingencies. Indeed, Group 2 mice were more proficient in a more challenging version of the task, where after leaving a depleted tower the next rewarding tower and direction were assigned randomly. Group 2 mice managed to maintain reward intake in this uncertain version, whereas Group 1 mice showed reduced reward gains, although there was considerable individual variability (some Group 2 mice performed poorly, and some Group 1 mice performed relatively well). This variability likely explains why, at the group level, there was no major difference in the exploitation of rewarded vs. unrewarded towers (Figures S9 and S10), although a trend for overexploitation of unrewarded towers seemed to develop in Group 1 mice. Still, a clear difference emerged: in the uncertain version, Group 2 mice markedly increased their number of runs between towers (RBT), whereas Group 1 mice did not (compare Figures S11D and S12D). This indicates that Group 2 mice were more explorative, most likely due to their prior experience with unrewarded outcomes when rewarded direction was reversed every morning session.

We acknowledge that the comparison between the two groups is not ideal, as Group 2 mice underwent longer training (this comparison was not initially planned). If anything, however, this strengthens our interpretation, since extended training under stable contingencies would likely have made Group 1 mice even more rigid. Altogether, our data reveal a form of meta-learning. Importantly, this meta-learning is not only characterized by improved efficiency within the trained task but also by a clear advantage in a novel protocol. Meta-learning is thought to arise from improved task representation (Samborska et al. 2022; Ambrogioni and Ólafsdóttir 2023; Hattori et al. 2023). Future work will investigate this possibility, although in our case we anticipate that it may simply result from an enhanced ability to update policies based on fewer evidence.

### Concluding remarks

In summary, the TFP offers a powerful framework for studying learning, flexible decision-making, and motor control. By training two groups of mice under distinct protocols, we showed that animals rapidly developed efficient strategies and adapted their choices and movements when confronted with sudden changes in foraging rules. Importantly, mice maintained a strong exploratory drive: they neither overexploited specific towers nor settled into stereotyped routines, which we interpret as evidence of naturalistic behavior. These experiments provide new insights into how reward influences motor control and uncovered a form of meta-learning, whose behavioral relevance was tested in alternative task versions, an approach, to our knowledge, rarely applied in rodents. The paradigm generates statistically robust datasets that comply with FAIR principles, and both hardware and software will be made openly available. Combined with neurophysiological and computational approaches, the TFP could also prove valuable for translational research, including the study of disease models, since foraging principles extend naturally to human behavior.

## Methods

### Experimental methods

#### Animals

Group 1 included 19 mice (9 males; 10 females) from two genetic backgrounds: C57BL/6N wildtype mice (Charles River, 4 mice) and C57BL/6J Eml1fl/fl and RhoAfl/fl “control” mice (Collins et al. 2019; Jackson et al. 2011, 8 and 7 mice, respectively). In these two transgenic lines, flox sequences were inserted around specific exons from the Eml1 or RhoA genes without interfering with the expression of these two genes. Eml1fl/fl and RhoAfl/fl “control” mice were therefore considered as “wild-type”. C57BL/6N wild type mice being used as controls for another study (to be published), these mice underwent anesthesia for 5 min (isoflurane 2.5-3%) every other day, after the end of the daily experimental sessions.

Group 2 included 32 C57BL/6J (20 males; 12 females) mice expressing Cre recombinase under the DRD1 or A2A promoter (DRD1-Cre or A2A-Cre, 8 and 7 mice, respectively, Gong et al. 2007; Durieux et al. 2009). Group 2 mice underwent a stereotaxic surgery under deep anesthesia (Isoflurane 2,5-3%) 1 week prior to the first familiarization session, to perform chemogenetic manipulations of striatal projection neurons (results not described in this study). All mice were 8 to 12 weeks old at the start of the experiments and were group-housed (2–3 per cage) in temperature-controlled, ventilated racks. A reversed 12 h light/dark cycle was maintained, and all behavioral experiments were conducted during the dark phase. Food was available ad libitum throughout the experiment. Mice underwent a controlled water restriction protocol. For this restriction, water bottles were removed one day prior to the beginning of the foraging sessions (i.e., after handling and familiarization sessions; see below). Body weight was monitored twice daily (before and after behavioral sessions), and animals were maintained at or above 85% of their baseline body weight throughout water restriction. All procedures were conducted in accordance with institutional and national ethical guidelines for animal research (European Community Directive 86/609/EEC) and were approved by the French Ministry of Higher Education and Research (Ministère de l’enseignement supérieur et de la recherche, France, Authorization #46285).

#### Task apparatus and foraging experiments

##### Apparatus

The behavioral task was conducted in a custom-designed square arena (83 × 83 cm) enclosed by opaque 9-cm-high walls (Figure 1). At the center of each quadrant, a square tower (12 × 12 cm wide, 9 cm high) was placed, resulting in four fixed tower locations: northwest, northeast, southwest, and southeast. Each of the four vertical faces of every tower was equipped with a protruding water port (total of 16 ports), consisting of a 0.9 mm diameter tube centered on the wall and positioned 2 cm above the arena floor. The tube extended 2 cm outward and was connected to an internal water reservoir within the tower, which was covered and inaccessible to the animal. Water drops (2–5 µL) were delivered via solenoid valves (VAC-20 PSIG, 12 V, Parker), each controlled independently through digital pulses, enabling precise and localized closed-loop water delivery. The floor of the arena was made of transparent Plexiglas and elevated 140 cm above the ground to allow for video tracking from below using an infrared-sensitive camera (Grasshopper3 GS3_U3_41C6NIR, Point Grey Research), operating at 25 Hz. The transparent Plexiglas floor was laser-cut into interlocking puzzle-shaped tiles to facilitate cleaning and modularity. Each tile included pre-drilled holes precisely positioned to accommodate vertical support pillars used to secure the tower walls and outer arena walls. This design allows for flexible reconfiguration of the arena layout, including change in the number, shape and position of the towers. All experiments were conducted in complete darkness. Upon publication, all the hardware components along with instruction to build the apparatus will be available in a public repository at https://github.com/robbe-neuroteam/TowersForagingPark-Hardware.

##### Closed-loop water delivery

Mouse position was tracked and video frames were processed in real time using a custom Python script using a background subtraction algorithm provided by the open-source OpenCV library. The extracted positional data was processed online and communicated to a Raspberry Pi 3B+ to control water delivery. The system employed a custom Raspberry Pi HAT integrating a 16-bit input/output expander (MCP23017) and two Darlington transistor arrays (ULN2803A) to independently trigger the 16 solenoid valves.

Four trapezoidal regions (trapezes) were defined around each tower, each containing a water port. A drop of water was delivered when the mouse crossed the boundary between two adjacent trapezes surrounding a given tower (depending on foraging rules). This crossing triggered (or not) a digital pulse to open the corresponding valve. Drops were delivered even if the mouse did not stop or lick at the port, allowing for passive accumulation of drops early in training sessions.

##### General behavioral procedure

Prior to behavioral testing, all the mice were first accustomed to the experimenter through two daily “handling” sessions of 5 to 10 minutes for 4-5 consecutive days. Then, they underwent two 12 minutes-long (Group 1) or 15 minutes-long (Group 2) familiarization sessions (one in the morning and one in the afternoon), during which mice could freely explore the arena. No water was delivered. Throughout the figures of this article, these familiarization sessions are referred to as sessions 1 and 2 (day 1). After performing these sessions, mice entered the water-restricted protocol. From day 2 onward, they were subjected to two daily foraging sessions lasting 12 minutes each, one in the morning and one in the afternoon, interspaced by at least 4 hours. The number of sessions and their duration allowed mice to maintain their weight at about 90 to 95% of their body weight before water restriction. Well-trained mice stopped foraging for water after 5 to 8 minutes, most likely because they started looking for solid food. Throughout the entire protocol, water was only accessible during the foraging sessions, except for mice whose weight dropped below 85% of their pre-restriction weight, in which case supplemental water was provided (this occurred only for a few mice at the beginning of training).

##### Experiment 1 Procedure

Starting on day 2, mice from Group 1 performed two daily 12 min-long foraging sessions (one in the morning and one in the afternoon, spaced by at least 4 hours), during which they could obtain drops of water by going from one water port to the next one. Critically, the mice had to adapt to the two following rules. First, only counterclockwise (CCW, from the camera viewpoint) turns around the towers triggered reward delivery at the next water port. Second, each time mice approached a tower, a random number of available drops was randomly assigned (between 4 and 12). Mice could leave a tower at any time, regardless of whether it was fully or partially depleted. After leaving a tower, rewards could only be obtained at the remaining three towers. The rewarded direction was maintained for 6 days (i.e., 12 CCW sessions) after which it was reversed to clockwise (CW) for three days (i.e., 6 CW sessions).

##### Experiment 2 Procedure

Starting on day 2, mice from Group 2 performed two daily 12-minute foraging sessions (one in the morning and one in the afternoon, spaced by at least 4 hours). On day 2, mice from Group 2 could obtain an unlimited number of rewards by performing either clockwise (CW) or counterclockwise (CCW) quadrant transitions (QTs), with both directions rewarded during both sessions. From day 3, the rewarded direction was fixed for the day and alternated each morning (e.g., if CCW was rewarded on day 3, CW was rewarded on day 4, and so on). From day 5 to day 10, the maximum number of rewards available per tower visit was randomly set between 4 and 12, as in Experiment 1. From day 11 to day 17, as part of a separate study, mice followed the same foraging protocol (two daily sessions with the rewarded direction changing each morning) but received an intraperitoneal injection of clozapine-N-oxide (CNO, 1 mg/kg) 30 minutes before the morning sessions of day 11 and day 16 (to perturbate the activity of striatal projection neurons). Data from days 11 to 17 are not included in this study, but the behavioral perturbations induced by CNO, which will be reported in a separate article, were fully reversed by the afternoon sessions. Indeed, comparing the main behavioral parameters (preference for rewarded vs non rewarded QTs, number QTs, speed of QTs, variability of QTs) the day before the first CNO injection (Figure 8, lasts 2 sessions) versus the day after the second injection (Figure 9, first 2 sessions) indicated very similar performance. Although we cannot exclude the possibility that these chemogenetic manipulations affected covert aspects of behavior, using the same animals across multiple protocols (rather than training an additional group of animals) satisfies ethical recommendations to limit the number of animals.

##### Experiment 3 Procedure

After adapting to the directional switch (once for Group 1 or across several days for Group 2), mice were challenged in a protocol in which both the rewarded direction (CW or CCW) and the identity of the rewarded tower (northwest, northeast, southeast, or southwest) were randomly reassigned each time a mouse left a fully depleted tower. The maximum number of rewards per tower visit was set as in Experiments 1 and 2 (i.e., randomly between 4 and 12). This procedure began on day 11 for Group 1 and day 18 for Group 2, lasting 3 and 4 days, respectively.

### Numerical methods

#### Data and code availability

Each experimental session generates two types of output files: the video of the session (saved in AVI format) and three CSV files containing the mouse’s positions, information related to trapeze switches and task parameters (key inputs of the acquisition code). A standardized electronic lab book is updated for every animal to store other important metadata, not directly necessary for data analysis (animal’s weight, age, sex, lineage and genotype, date of birth, name of experimenter and general protocol). The data processing pipeline was implemented using custom Python scripts. All experimental data (CSV files) and Jupyter notebooks used for data processing, behavioral analysis, and figure generation will be made publicly available upon publication of the manuscript.

Our code and data comply with the Findability, Accessibility, Interoperability, and Reusability (FAIR) principles.

(F): Upon publication, our data and codes to generate the figures will be deposited in stable public repositories: OSF (https://osf.io/23f76/) for the data and GitHub (https://github.com/robbe-neuroteam/4TowersTask_MethodPaper) for the code. The data are extensively described in a dedicated file on OSF.

(A): Both the data and code are open-source.

(I): The code is interoperable, as it was developed exclusively in Python 3.12 and can therefore be used on any computer and operating system that supports this version. Data files follow a clear nomenclature, and the code uses explicit variable names and is extensively commented, which facilitates reuse. Applying the code to new datasets requires minimal manual intervention. (R): The data and code are free to use provided that this article is cited. The origins of the data sets are clearly stated.

#### Data processing

Behavioral data were saved as CSV files generated by the acquisition software (see above). Three primary data sources were used: (1) time-stamped x/y coordinates of the animal’s centroid, (2) information each time the animal just crossed the boundary between two adjacent trapezes (e.g., time of this crossing, cardinal identity of the tower [NW, NE, SW, or SE], cardinal identity of the trapezes where the animal is [N, E, S, or W], cardinal identity of the trapezes where the animal was on the previous video frame, information related to the whether this trapeze switch will deliver a reward) and (3) session metadata. The first 15 seconds of each session were excluded to remove tracking inaccuracies due to background subtraction initialization. Additionally, positional data were filtered to retain only values within a 1–500 pixel range, discarding outliers. Spatial coordinates were converted from pixels to centimeters using a scaling factor derived from the physical dimensions of the arena. Positions were smoothed using a 1D Gaussian filter (σ = 1), and these smoothed trajectories were used to compute distances, instantaneous speeds, and angular velocities (in degrees per second).

Running epochs were automatically identified based on a speed threshold (> 7 cm/s), with a minimum duration of 0.3 s and tolerance for short pauses (< 0.1 s). To refine the temporal boundaries of each running epoch, the initial and final time points were further adjusted based on local acceleration dynamics. Specifically, for the onset of each run, the algorithm traced backward from the speed threshold-crossing point to identify the earliest time point at which acceleration dropped below 10% of the acceleration observed at the threshold. This allowed inclusion of pre-threshold data points that approximated better movement initiation. A similar procedure was applied at the end of the run, tracing forward from the point where speed dropped below the threshold to detect when deceleration fell below 10% of the prior peak, providing a better approximation of the moment when locomotor activity ceased. This correction ensured that each epoch more accurately reflected the full extent of runs rather than relying solely on fixed speed thresholds.

Each running epoch was classified based on the animal’s position at the beginning and end of the detected run. Runs were categorized as exploratory runs, approaches toward a tower, runs between towers (RBTs), or runs around a tower. In this study, we focused on RBTs and runs around a tower. RBTs were defined as runs starting and ending in trapezes located around two different towers (i.e., when mice run from one water port of a given tower to a water port located around a different tower). Runs around a tower were defined as runs starting and ending in trapezes located around the same tower, with the mice crossing at least one border between two contiguous trapezes. Almost all runs around a tower consisted of quarter-turns (QTs), in which mice moved from one water port to the next (either in a clockwise or counterclockwise direction). A very small number of runs around a tower corresponded to half-turns (e.g., a run from the south water port to the north water port of a given tower without stopping at the east or west port). In extremely rare cases, mice started and finished their run around the same tower but after a long detour. To avoid comparing heterogeneous turns around a tower, we restricted our analysis to QTs. Practically, QTs were runs around a tower during which mice performed only one trapeze switch, covered a distance shorter than 30 cm, and had a duration shorter than 2 seconds.

#### Foraging analyses

To quantify the extent of mice engagement in reward collection (Figures 2 and S4), we computed their time spent within different zones of the arena (near the peripheral walls vs in the trapezes surrounding the towers).

To quantify how much mice exploit a given tower, we then defined visits as sequences of QTs performed around a tower (Figures 5 and S9). A QT is considered as the start of a visit if the previous run epoch was not a QT in the same tower. For each visit, we computed the number of QTs, the number of rewarded QTs, the maximum number of rewards available and the starting time of the visit.

#### Kinematic analyses

Basic kinematic parameters (duration, distance traveled, mean and peak speed) were extracted for all runs (Figures 3, 6, 8, 9, S2 and S3). For QTs, additional features were extracted: direction (CW or CCW), and reward status (rewarded or not) (Figures 3, 5, 6, 7, 8, 9, S1, S2, S6, S7 and S8). Concerning Group 2 mice (those for which the rewarded direction alternated every day), most of the animals started with CW QTs being rewarded on day 3 while a subgroup started with CCW QTs being rewarded. To examine whether, independently from the rewarded direction on day 3, all these mice updated across sessions their preference for rewarded and unrewarded QTs (Figure 8), we arbitrarily relabeled CCW QTs as CW in the small subgroup for which CCW QTs were rewarded on day 3.

To evaluate the similarity between QTs trajectories, we computed the Fréchet distance, which is the minimum cord-length needed to connect a point moving along one trajectory to a point moving along another trajectory. Fréchet distance was computed across all possible pairs of QTs performed during a given session, separately for CW and CCW QTs (Figures 3, 6, 8, 9 and S1). This involved translating and rotating trajectories relative to a single point (a single corner of a single tower). First, the coordinates of the positions forming each QT (initially expressed in the arena reference frame) are transformed to be expressed relative to the specific tower corner around which the QT occurs. This step ensures that all QTs share a common reference point Then, QT trajectories around the SW, NW and NE corners are rotated CCW by 90, 180 and 270 degrees, respectively. Thus, all the CW and CCW QTs performed during a session around different corners of different towers became aligned, allowing the quantification of Fréchet distances between pairs of CCW QTs and pairs of CW QTs. We use the median of all the Fréchet distances computed from all the pairs of CW and CCW QT as measure of the variability in their trajectory.

#### Statistical analyses

In all the figures showing performance metrics across sessions and mice, individual data (thin colored lines), group median (thick lines) and interquartile range (error bars) are represented (Figures 2, 3, 4, 5, 6, 8, 9, S1, S2, S4, S5, S8, S9, S10). The number of mice in each group is indicated in the figure or in the figure legend (Group 1 n= 19, Group 2 n=32).

To assess differences in speed or Fréchet distance between CW and CCW QTs between conditions across sessions, we applied non-parametric paired-label permutation tests (Figures S1 and S2). For each session, the metric of interest was extracted for each mouse in each group, and a summary statistic was computed per group. The observed difference between groups was then compared to a null distribution obtained by randomly swapping condition labels within mice across a fixed number of permutations. For Figures S1B and S2B, we computed group medians and used 500 permutations to generate the null distribution of median differences. To verify that the low values of speed (or high value of Fréchet distance) of CW QTs was not an artefact related to their very small number, we randomly downsampled for each animal and each session, a smaller number of CCW QTs such as it matches the number of CW QTs performed. Then we recomputed the median speed and Fréchet distance for this small number of CCW QTs. We repeated the downsampling 100 times. The red shaded areas in Figure S1C and S2C are the interquartile distribution of these recomputed speeds and Fréchet distances. The original values of CCW speeds and Fréchet distances are always contained inside this interquartile interval, showing that the small number of CW QTs are not responsible for small speed values nor high values of Fréchet distance.

For Figure S8ABC, to quantify the difference between Group 1 and 2 on several metrics, we computed group means and used 500 permutations with a two-sided test per session. Session-wise *p*-values were calculated as the proportion of permutations yielding a difference at least as extreme as the observed one. Confidence intervals (typically 90%) were derived from the null distribution, and significant sessions (*p* < 0.05) were marked in the visualizations. This approach avoids distributional assumptions and accounts for within-subject variability across time.

To assess within-session evolution in turn speed, we focused on CCW turn speeds (Figure S3A). For each session containing at least 100 CCW trials, we computed the slope of the linear regression between trial rank (i.e., position within the session) and maximal turn speed. A one-sided permutation test was performed to assess whether the observed slope was significantly more negative than expected under the null hypothesis of no temporal structure. At each iteration (n = 10000), CCW turn speeds were randomly shuffled, and the regression slope was recomputed to generate a null distribution. The p-value was calculated as the proportion of permuted slopes less than or equal to the observed slope. The evolution of slope values across sessions is shown for each mouse (Figure S3B). Sessions with significant negative slopes (*p < 0.05*) are marked with circles, non-significant ones with crosses. The same analysis was applied to the Pearson correlation coefficient (r) between trial rank and turn speed, highlighting sessions with significant effects (Figure S3C). For each mouse, we recorded the number of sessions showing a significant negative slope (*p < 0.05*), summarizing the prevalence of within-session effects at the individual level (Figure S3D).

To compare paired data (Figure 2E, S4E), we used a paired Wilcoxon test with a significance criterion of *p < 0.05*.

In figure 5C, we quantified tower preference by comparing the time mice spent around each tower to the chance level of 0.25 (equal time at each of the 4 towers). For each mouse, tower and session, we computed the proportion of time spent near that tower, resulting in a distribution of ratios. To assess whether mice preferred certain towers, we compared the median of each distribution to the chance level using a one-sample Wilcoxon test (after subtracting 0.25 from each value), with a significance criterion of *p < 0.05*.

## Acknowledgments

This work has been supported by the French National Research Agency ANR-20-CE16-0002/Corticostriatal; from the “Investissements d’Avenir” French Government program managed by the French National Research Agency (ANR-16-CONV-0001), Initiative d’Excellence d’Aix-Marseille Université, AMX-22-RE-AB-007/Neuradventure; the Ministère de l’Enseignement Supérieur et de la Recherche (KA, AF), a neuroschool PhD fellowship (Initiative d’Excellence d’Aix-Marseille Université, AMX-19-IET-004 and ANR-17-EURE-0029; KA, AF),, from the Fondation pour la Recherche Médicale (FDT202204014828; TM).

We thank Ingrid Bureau, Ahmed El Haddy, Chirstophe Eloy and Timothy Verstynen for discussions at different stages of the development of this task. Nihaad Paraouty, Alexy Louis, Raed Ben Yedder, Alice Le Bar, Remi Proville for technical help at an early stage of the task development. We thank the members of the Corticio-Basal Ganglia Circuit Team and INMED’s animal facility. We thank Rosa Cossart, Ahmed El Haddy, Christophe Eloy, Jerome Epsztein, Olivier Manzoni, Sarah Mondoloni, and Zelda Timmel for their helpful feedback on the manuscript. We thank Ahmed El Haddy for suggesting the name of the task.

## References

Addicott, M. A., J. M. Pearson, M. M. Sweitzer, D. L. Barack, and M. L. Platt. 2017. “A Primer on Foraging and the Explore/Exploit Trade-Off for Psychiatry Research.” Neuropsychopharmacology 42 (10): 1931–39. 10.1038/npp.2017.108.

Aldridge, J. Wayne, and Kent C. Berridge. 1998. “Coding of Serial Order by Neostriatal Neurons: A ‘Natural Action’ Approach to Movement Sequence.” ARTICLE. Journal of Neuroscience 18 (7): 2777–87. 10.1523/JNEUROSCI.18-07-02777.1998.

Ambrogioni, Luca, and H. Freyja Ólafsdóttir. 2023. “Rethinking the Hippocampal Cognitive Map as a Meta-Learning Computational Module.” Trends in Cognitive Sciences 27 (8): 702–12. 10.1016/j.tics.2023.05.011.

Barack, David L. 2024. “What Is Foraging?” Biology & Philosophy 39 (1): 3. 10.1007/s10539-024-09939-z.

Barack, David L., Vera U. Ludwig, Felipe Parodi, et al. 2024. “Attention Deficits Linked with Proclivity to Explore While Foraging.” Proceedings of the Royal Society B: Biological Sciences 291 (2017): 20222584. 10.1098/rspb.2022.2584.

Basso, Michele A., and Robert H. Wurtz. 1997. “Modulation of Neuronal Activity by Target Uncertainty.” Nature 389 (6646): 66–69. 10.1038/37975.

Berridge, K. C., J. C. Fentress, and H. Parr. 1987. “Natural Syntax Rules Control Action Sequence of Rats.” Behavioural Brain Research 23 (1): 59–68. 10.1016/0166-4328(87)90242-7.

Britten, K. H., M. N. Shadlen, W. T. Newsome, and J. A. Movshon. 1992. “The Analysis of Visual Motion: A Comparison of Neuronal and Psychophysical Performance.” Articles. Journal of Neuroscience 12 (12): 4745–65. 10.1523/JNEUROSCI.12-12-04745.1992.

Brunton, Bingni W., Matthew M. Botvinick, and Carlos D. Brody. 2013. “Rats and Humans Can Optimally Accumulate Evidence for Decision-Making.” Science 340 (6128): 95–98. 10.1126/science.1233912.

Burgat, Florence, ed. 2010. Penser le comportement animal. In Penser le comportement animal. Natures sociales. Éditions de la Maison des sciences de l’homme. https://books.openedition.org/editionsmsh/12864.

Canaveral, Cesar Augusto, William Lata, Andrea M. Green, and Paul Cisek. 2024. “Biomechanical Costs Influence Decisions Made during Ongoing Actions.” Journal of Neurophysiology 132 (2): 461–69. 10.1152/jn.00090.2024.

Carland, Matthew A., David Thura, and Paul Cisek. 2019. “The Urge to Decide and Act: Implications for Brain Function and Dysfunction.” The Neuroscientist 25 (5): 491–511. 10.1177/1073858419841553.

Charnov, Eric L. 1976. “Optimal Foraging, the Marginal Value Theorem.” Theoretical Population Biology 9 (2): 129–36. 10.1016/0040-5809(76)90040-X.

Chen, Xiuli, Peter Holland, and Joseph M Galea. 2018. “The Effects of Reward and Punishment on Motor Skill Learning.” *Current Opinion in Behavioral Sciences*, Habits and Skills, vol. 20 (April): 83–88. 10.1016/j.cobeha.2017.11.011.

Chirimuuta, M. 2024. *The Brain Abstracted: Simplification in the History and Philosophy of Neuroscience*. MIT Press. https://mitpress.mit.edu/9780262548045/the-brain-abstracted/.

Chrzanowska, Anna, Mateusz Kostecki, Natalia Krasilshchikova, et al. 2025. “Rethinking Naturalistic Behavior: The Terzolas Manifesto.” Preprint, OSF, February 25. 10.31219/osf.io/ch3an_v1.

Cinotti, François, Virginie Fresno, Nassim Aklil, et al. 2019. “Dopamine Blockade Impairs the Exploration-Exploitation Trade-off in Rats.” Scientific Reports 9 (1): 6770. 10.1038/s41598-019-43245-z.

Cisek, Paul, and Andrea M. Green. 2024. “Toward a Neuroscience of Natural Behavior.” Current Opinion in Neurobiology 86: 102859.

Cisek, Paul, and John F. Kalaska. 2010. “Neural Mechanisms for Interacting with a World Full of Action Choices.” Annual Review of Neuroscience 33 (1): 269–98. 10.1146/annurev.neuro.051508.135409.

Cisek, Paul, and Alexandre Pastor-Bernier. 2014. “On the Challenges and Mechanisms of Embodied Decisions.” Philosophical Transactions of the Royal Society B: Biological Sciences 369 (1655): 20130479. 10.1098/rstb.2013.0479.

Codol, Olivier, Peter J. Holland, Sanjay G. Manohar, and Joseph M. Galea. 2020. “Reward-Based Improvements in Motor Control Are Driven by Multiple Error-Reducing Mechanisms.” The Journal of Neuroscience: The Official Journal of the Society for Neuroscience 40 (18): 3604–20. 10.1523/JNEUROSCI.2646-19.2020.

Collins, Stephan C., Ana Uzquiano, Mohammed Selloum, et al. 2019. “The Neuroanatomy of Eml1 Knockout Mice, a Model of Subcortical Heterotopia.” Journal of Anatomy 235 (3): 637–50. 10.1111/joa.13013.

Cos, Ignasi, Nicolas Bélanger, and Paul Cisek. 2011. “The Influence of Predicted Arm Biomechanics on Decision Making.” Journal of Neurophysiology 105 (6): 3022–33. 10.1152/jn.00975.2010.

Cowen, Stephen L., and Bruce L. McNaughton. 2007. “Selective Delay Activity in the Medial Prefrontal Cortex of the Rat: Contribution of Sensorimotor Information and Contingency.” Journal of Neurophysiology 98 (1): 303–16. 10.1152/jn.00150.2007.

De Chaumont, Fabrice, Renata Dos-Santos Coura, Pierre Serreau, et al. 2012. “Computerized Video Analysis of Social Interactions in Mice.” Nature Methods 9 (4): 410–17.

Dennis, Emily Jane, Ahmed El Hady, Angie Michaiel, et al. 2021. “Systems Neuroscience of Natural Behaviors in Rodents.” Symposium and Mini-Symposium. Journal of Neuroscience 41 (5): 911–19. 10.1523/JNEUROSCI.1877-20.2020.

Despret, Vinciane. 2015. “Thinking Like a Rat.” Angelaki 20 (2): 121–34. 10.1080/0969725X.2015.1039849.

Dhawale, Ashesh K., Yohsuke R. Miyamoto, Maurice A. Smith, and Bence P. Ölveczky. 2019. “Adaptive Regulation of Motor Variability.” Current Biology 29 (21): 3551–3562.e7. 10.1016/j.cub.2019.08.052.

Dhawale, Ashesh K., Maurice A. Smith, and Bence P. Ölveczky. 2017. “The Role of Variability in Motor Learning.” Annual Review of Neuroscience 40 (Volume 40, 2017): 479–98. 10.1146/annurev-neuro-072116-031548.

Durieux, Pierre F., Bertrand Bearzatto, Stefania Guiducci, et al. 2009. “D2R Striatopallidal Neurons Inhibit Both Locomotor and Drug Reward Processes.” Nature Neuroscience 12 (4): 393–95. 10.1038/nn.2286.

El Hady, Ahmed, Jacob D. Davidson, and Deborah M. Gordon. 2019. “Editorial: An Ecological Perspective on Decision-Making: Empirical and Theoretical Studies in Natural and Natural-Like Environments.” Frontiers in Ecology and Evolution 7 (December). 10.3389/fevo.2019.00461.

Euston, David R., and Bruce L. McNaughton. 2006. “Apparent Encoding of Sequential Context in Rat Medial Prefrontal Cortex Is Accounted for by Behavioral Variability.” Articles. Journal of Neuroscience 26 (51): 13143–55. 10.1523/JNEUROSCI.3803-06.2006.

Fentress, J. C., and F. P. Stilwell. 1973. “Letter: Grammar of a Movement Sequence in Inbred Mice.” Nature 244 (5410): 52–53. 10.1038/244052a0.

Georgopoulos, Apostolos P., Andrew B. Schwartz, and Ronald E. Kettner. 1986. “Neuronal Population Coding of Movement Direction.” Science 233 (4771): 1416–19. 10.1126/science.3749885.

Gomez-Marin, Alex, and Asif A. Ghazanfar. 2019. “The Life of Behavior.” Neuron 104 (1): 25–36. 10.1016/j.neuron.2019.09.017.

Gong, Shiaoching, Martin Doughty, Carroll R. Harbaugh, et al. 2007. “Targeting Cre Recombinase to Specific Neuron Populations with Bacterial Artificial Chromosome Constructs.” The Journal of Neuroscience: The Official Journal of the Society for Neuroscience 27 (37): 9817–23. 10.1523/JNEUROSCI.2707-07.2007.

Gouvea, Thiago S., Tiago Monteiro, Sofia Soares, Bassam V. Atallah, and Joseph J. Paton. 2014. “Ongoing Behavior Predicts Perceptual Report of Interval Duration.” Frontiers in Neurorobotics 8 (March). 10.3389/fnbot.2014.00010.

Hamid, Arif A., Jeffrey R. Pettibone, Omar S. Mabrouk, et al. 2016. “Mesolimbic Dopamine Signals the Value of Work.” Nature Neuroscience 19 (1): 1. 10.1038/nn.4173.

Hasnain, Munib A., Jaclyn E. Birnbaum, Juan Luis Ugarte Nunez, Emma K. Hartman, Chandramouli Chandrasekaran, and Michael N. Economo. 2025. “Separating Cognitive and Motor Processes in the Behaving Mouse.” Nature Neuroscience 28 (3): 640–53. 10.1038/s41593-024-01859-1.

Hattori, Ryoma, Nathan G. Hedrick, Anant Jain, et al. 2023. “Meta-Reinforcement Learning via Orbitofrontal Cortex.” Nature Neuroscience 26 (12): 2182–91. 10.1038/s41593-023-01485-3.

Hayden, Benjamin Y., John M. Pearson, and Michael L. Platt. 2011. “Neuronal Basis of Sequential Foraging Decisions in a Patchy Environment.” Nature Neuroscience 14 (7): 933–39. 10.1038/nn.2856.

Jackson, Ben, Karine Peyrollier, Esben Pedersen, et al. 2011. “RhoA Is Dispensable for Skin Development, but Crucial for Contraction and Directed Migration of Keratinocytes.” *Molecular Biology of the Cell*, ahead of print, January 5. world. 10.1091/mbc.e09-10-0859.

Jami, Ali, Sajjad Abbaszade, and Abdol-Hossein Vahabie. 2025. “A Review on Exploration–Exploitation Trade-off in Psychiatric Disorders.” BMC Psychiatry 25 (1): 420. 10.1186/s12888-025-06837-w.

Jin, Xin, and Rui M. Costa. 2010. “Start/Stop Signals Emerge in Nigrostriatal Circuits during Sequence Learning.” Nature 466 (7305): 457–62. 10.1038/nature09263.

Jurado-Parras, Maria-Teresa, Mostafa Safaie, Stefania Sarno, et al. 2020. “The Dorsal Striatum Energizes Motor Routines.” Current Biology 30 (22): 4362–4372.e6. 10.1016/j.cub.2020.08.049.

Kawai, Risa, Timothy Markman, Rajesh Poddar, et al. 2015. “Motor Cortex Is Required for Learning but Not for Executing a Motor Skill.” Neuron 86 (3): 800–812. 10.1016/j.neuron.2015.03.024.

Killeen, Peter R., and J. Gregor Fetterman. 1988. “A Behavioral Theory of Timing.” Psychological Review 95 (2): 274.

Kilpatrick, Zachary P., Jacob D. Davidson, and Ahmed El Hady. 2021a. “Uncertainty Drives Deviations in Normative Foraging Decision Strategies.” Journal of The Royal Society Interface 18 (180): 20210337. 10.1098/rsif.2021.0337.

Kilpatrick, Zachary P., Jacob D. Davidson, and Ahmed El Hady. 2021b. “Uncertainty Drives Deviations in Normative Foraging Decision Strategies.” Journal of The Royal Society Interface 18 (180): 20210337. 10.1098/rsif.2021.0337.

Krakauer, John W., Asif A. Ghazanfar, Alex Gomez-Marin, Malcolm A. MacIver, and David Poeppel. 2017. “Neuroscience Needs Behavior: Correcting a Reductionist Bias.” Neuron 93 (3): 480–90. 10.1016/j.neuron.2016.12.041.

Krakauer, John W., Alkis M. Hadjiosif, Jing Xu, Aaron L. Wong, and Adrian M. Haith. 2019. “Motor Learning.” In Comprehensive Physiology. American Cancer Society. 10.1002/cphy.c170043.

Lemke, Stefan M., Dhakshin S. Ramanathan, Ling Guo, Seok Joon Won, and Karunesh Ganguly. 2019. “Emergent Modular Neural Control Drives Coordinated Motor Actions.” Nature Neuroscience 22 (7): 1122–31. 10.1038/s41593-019-0407-2.

Leppänen, Pia K., Niklas Ravaja, and S. B. M. Ewalds-Kvist. 2006. “Twenty-Three Generations of Mice Bidirectionally Selected for Open-Field Thigmotaxis: Selection Response and Repeated Exposure to the Open Field.” Behavioural Processes 72 (1): 23–31.

Manohar, Sanjay G., Trevor T.-J. Chong, Matthew A. J. Apps, et al. 2015. “Reward Pays the Cost of Noise Reduction in Motor and Cognitive Control.” Current Biology 25 (13): 1707–16. 10.1016/j.cub.2015.05.038.

Manohar, Sanjay G., Kinan Muhammed, Sean J. Fallon, and Masud Husain. 2019. “Motivation Dynamically Increases Noise Resistance by Internal Feedback during Movement.” Neuropsychologia 123 (February): 19–29. 10.1016/j.neuropsychologia.2018.07.011.

Markowitz, Jeffrey E., Winthrop F. Gillis, Celia C. Beron, et al. 2018. “The Striatum Organizes 3D Behavior via Moment-to-Moment Action Selection.” Cell 174 (1): 44–58.e17. 10.1016/j.cell.2018.04.019.

Markowitz, Jeffrey E., Winthrop F. Gillis, Maya Jay, et al. 2023. “Spontaneous Behaviour Is Structured by Reinforcement without Explicit Reward.” Nature 614 (7946): 7946. 10.1038/s41586-022-05611-2.

Mathis, Alexander, Pranav Mamidanna, Kevin M. Cury, et al. 2018. “DeepLabCut: Markerless Pose Estimation of User-Defined Body Parts with Deep Learning.” Nature Neuroscience 21 (9): 1281–89.

Mizes, Kevin G. C., Jack Lindsey, G. Sean Escola, and Bence P. Ölveczky. 2022. *Similar Striatal Activity Exerts Different Control over Automatic and Flexible Motor Sequences*. Preprint. Neuroscience. 10.1101/2022.06.13.495989.

Mohebi, Ali, Jeffrey R. Pettibone, Arif A. Hamid, et al. 2019. “Dissociable Dopamine Dynamics for Learning and Motivation.” Nature 570 (7759): 7759. 10.1038/s41586-019-1235-y.

Morvan, Thomas, Christophe Eloy, and David Robbe. 2024. “Running, Fast and Slow: The Dorsal Striatum Sets the Cost of Movement During Foraging.” Preprint, bioRxiv, June 1. 10.1101/2024.05.31.596850.

Musall, Simon, Matthew T. Kaufman, Ashley L. Juavinett, Steven Gluf, and Anne K. Churchland. 2019. “Single-Trial Neural Dynamics Are Dominated by Richly Varied Movements.” Nature Neuroscience 22 (10): 1677–86. 10.1038/s41593-019-0502-4.

Oesch, Lukas T., Michael B. Ryan, and Anne K. Churchland. 2024. “From Innate to Instructed: A New Look at Perceptual Decision-Making.” Current Opinion in Neurobiology 86 (June): 102871. 10.1016/j.conb.2024.102871.

O’Keefe, John. 1979. “A Review of the Hippocampal Place Cells.” Progress in Neurobiology 13 (4): 419–39. 10.1016/0301-0082(79)90005-4.

Pekny, Sarah E., Jun Izawa, and Reza Shadmehr. 2015. “Reward-Dependent Modulation of Movement Variability.” Articles. Journal of Neuroscience 35 (9): 4015–24. 10.1523/JNEUROSCI.3244-14.2015.

Pereira, Talmo D., Nathaniel Tabris, Arie Matsliah, et al. 2022. “SLEAP: A Deep Learning System for Multi-Animal Pose Tracking.” Nature Methods 19 (4): 486–95. 10.1038/s41592-022-01426-1.

Pfeifer, Rolf, and Josh Bongard. 2006. *How the Body Shapes the Way We Think: A New View of Intelligence*. The MIT Press. 10.7551/mitpress/3585.001.0001.

Ray, Saikat, Itay Yona, Nadav Elami, et al. 2025. “Hippocampal Coding of Identity, Sex, Hierarchy, and Affiliation in a Social Group of Wild Fruit Bats.” Science (New York, N.Y.) 387 (6733): eadk9385. 10.1126/science.adk9385.

Richelle, and Lejeune. 1980. *Time in Animal Behaviour*. Elsevier. 10.1016/C2009-0-00224-4.

Robbe, David. 2023. “Lost in Time: Relocating the Perception of Duration Outside the Brain.” Neuroscience & Biobehavioral Reviews 153 (October): 105312. 10.1016/j.neubiorev.2023.105312.

Romo, Ranulfo, Carlos D. Brody, Adrián Hernández, and Luis Lemus. 1999. “Neuronal Correlates of Parametric Working Memory in the Prefrontal Cortex.” Nature 399 (6735): 470–73.

Rosenberg, Matthew, Tony Zhang, Pietro Perona, and Markus Meister. 2021. “Mice in a Labyrinth Show Rapid Learning, Sudden Insight, and Efficient Exploration.” eLife 10 (July): e66175. 10.7554/eLife.66175.

Rueda-Orozco, Pavel E, and David Robbe. 2015. “The Striatum Multiplexes Contextual and Kinematic Information to Constrain Motor Habits Execution.” Nature Neuroscience 18 (3): 453–60. 10.1038/nn.3924.

Safaie, Mostafa, Maria-Teresa Jurado-Parras, Stefania Sarno, et al. 2020. “Turning the Body into a Clock: Accurate Timing Is Facilitated by Simple Stereotyped Interactions with the Environment.” Proceedings of the National Academy of Sciences 117 (23): 13084–93. 10.1073/pnas.1921226117.

Sales-Carbonell, Carola, Wahiba Taouali, Loubna Khalki, et al. 2018. “No Discrete Start/Stop Signals in the Dorsal Striatum of Mice Performing a Learned Action.” Current Biology 28 (19): 3044–3055.e5. 10.1016/j.cub.2018.07.038.

Samborska, Veronika, James L. Butler, Mark E. Walton, Timothy E. J. Behrens, and Thomas Akam. 2022. “Complementary Task Representations in Hippocampus and Prefrontal Cortex for Generalizing the Structure of Problems.” Nature Neuroscience 25 (10): 1314–26. 10.1038/s41593-022-01149-8.

Sarel, Ayelet, Shaked Palgi, Dan Blum, Johnatan Aljadeff, Liora Las, and Nachum Ulanovsky. 2022. “Natural Switches in Behaviour Rapidly Modulate Hippocampal Coding.” Nature 609 (7925): 119–27. 10.1038/s41586-022-05112-2.

Shadmehr, Reza, and John W. Krakauer. 2008. “A Computational Neuroanatomy for Motor Control.” Experimental Brain Research 185 (3): 359–81. 10.1007/s00221-008-1280-5.

Shadmehr, Reza, Thomas R. Reppert, Erik M. Summerside, Tehrim Yoon, and Alaa A. Ahmed. 2019. “Movement Vigor as a Reflection of Subjective Economic Utility.” Trends in Neurosciences 42 (5): 323–36.

Sridhar, Gautam, Massimo Vergassola, João C. Marques, Michael B. Orger, Antonio Carlos Costa, and Claire Wyart. 2024. “Uncovering Multiscale Structure in the Variability of Larval Zebrafish Navigation.” Proceedings of the National Academy of Sciences 121 (47): e2410254121. world. 10.1073/pnas.2410254121.

Stephens, David W., and J. R. Krebs. 1986. *Foraging Theory*. Monographs in Behavior and Ecology. Princeton University Press.

Stringer, Carsen, Marius Pachitariu, Nicholas Steinmetz, Charu Bai Reddy, Matteo Carandini, and Kenneth D. Harris. 2019. “Spontaneous Behaviors Drive Multidimensional, Brainwide Activity.” Science 364 (6437): eaav7893. 10.1126/science.aav7893.

Summerside, Erik M., Reza Shadmehr, and Alaa A. Ahmed. 2018. “Vigor of Reaching Movements: Reward Discounts the Cost of Effort.” Journal of Neurophysiology 119 (6): 2347–57. 10.1152/jn.00872.2017.

Tai, Lung-Hao, A Moses Lee, Nora Benavidez, Antonello Bonci, and Linda Wilbrecht. 2012. “Transient Stimulation of Distinct Subpopulations of Striatal Neurons Mimics Changes in Action Value.” Nature Neuroscience 15 (9): 1281–89. 10.1038/nn.3188.

Takikawa, Yoriko, Reiko Kawagoe, Hideaki Itoh, Hiroyuki Nakahara, and Okihide Hikosaka. 2002. “Modulation of Saccadic Eye Movements by Predicted Reward Outcome.” Experimental Brain Research 142 (2): 284–91. 10.1007/s00221-001-0928-1.

The International Brain Laboratory, Valeria Aguillon-Rodriguez, Dora Angelaki, et al. 2021. “Standardized and Reproducible Measurement of Decision-Making in Mice.” eLife 10 (May): e63711. 10.7554/eLife.63711.

Torquet, N., F. Marti, C. Campart, et al. 2018. “Social Interactions Impact on the Dopaminergic System and Drive Individuality.” Nature Communications 9 (1): 3081. 10.1038/s41467-018-05526-5.

Ulanovsky, Nachum. 2025. *Natural Neuroscience: Toward a Systems Neuroscience of Natural Behaviors*. MIT Press.

Wang, Siyu, Blake Gerken, Julia R. Wieland, Robert C. Wilson, and Jean-Marc Fellous. 2023. “The Effects of Time Horizon and Guided Choices on Explore–Exploit Decisions in Rodents.” Behavioral Neuroscience 137 (2): 127–42. 10.1037/bne0000549.

Wiltschko, Alexander B., Tatsuya Tsukahara, Ayman Zeine, et al. 2020. “Revealing the Structure of Pharmacobehavioral Space through Motion Sequencing.” Nature Neuroscience 23 (11): 1433–43. 10.1038/s41593-020-00706-3.

Wystrach, Antoine. 2021. “Movements, Embodiment and the Emergence of Decisions. Insights from Insect Navigation.” Biochemical and Biophysical Research Communications 564 (July): 70–77. 10.1016/j.bbrc.2021.04.114.

Yoon, Tehrim, Robert B. Geary, Alaa A. Ahmed, and Reza Shadmehr. 2018. “Control of Movement Vigor and Decision Making during Foraging.” Proceedings of the National Academy of Sciences 115 (44). 10.1073/pnas.1812979115.

