## Supplementary Figures S1-S12 for "A Novel Foraging Task Reveals Cognitive and Motor Processes Underlying Behavioral Flexibility"

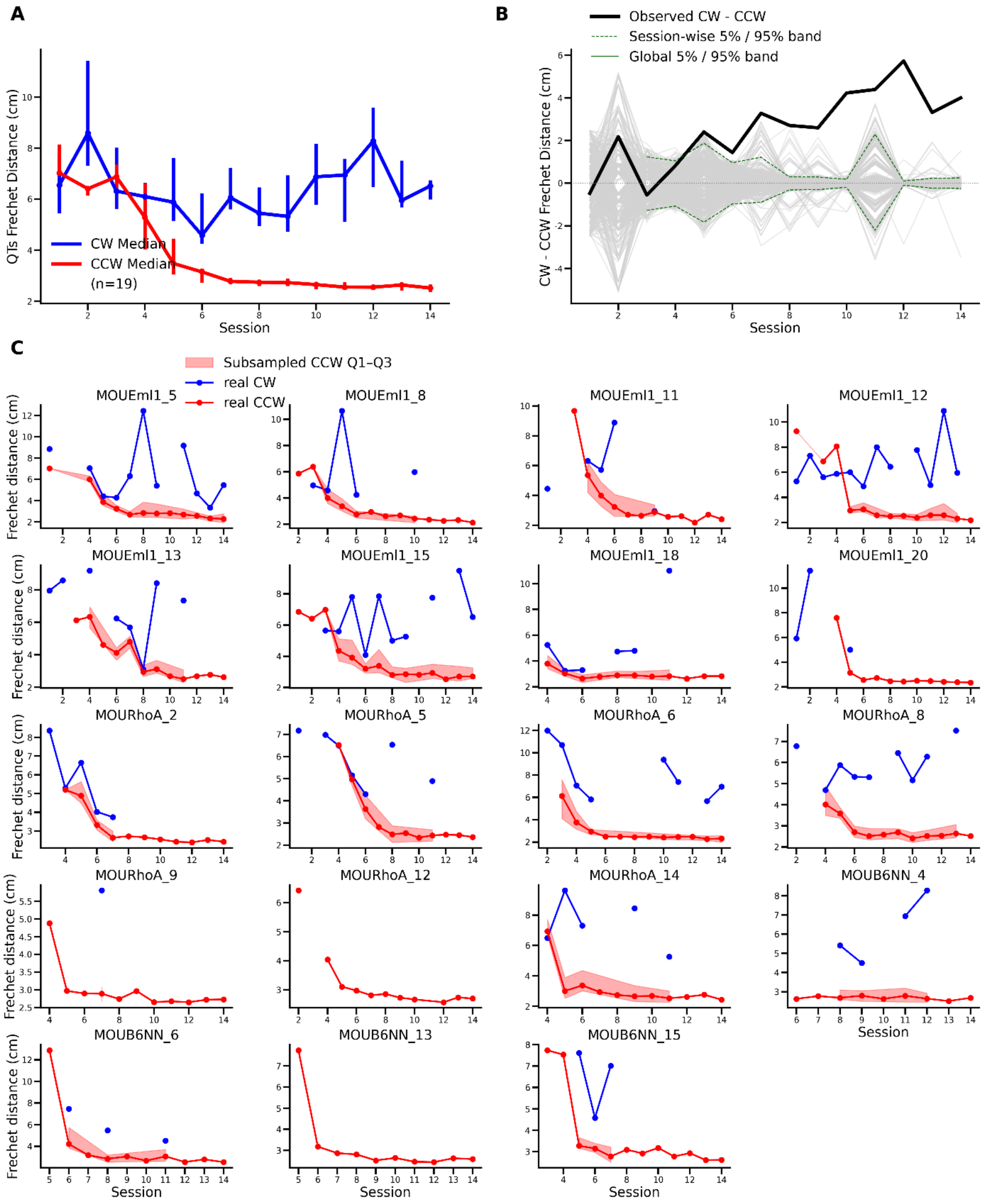


**Figure S1. Comparison of trajectory similarity between CCW and CW** QTs **across sessions. A)** Evolution of median Fréchet distances between QTs, computed separately for CCW (rewarded direction, red) and CW (unrewarded direction, blue) turns. **B)** Permutation test (n=500 iterations) comparing Fréchet distances of CW and CCW QTs. The solid black line shows the observed difference (CW minus CCW) in median Fréchet distance for each session. Dashed green lines indicate the global 5% and 95% confidence bands computed from the null distribution obtained by shuffling trajectory labels within each session. Solid dark green lines mark the session-wise thresholds corresponding to a z-score of ±2. Observed differences falling outside the dashed green confidence band indicate statistically significant divergence between CW and CCW QT similarity over time. **C)** Same analysis as in panel A, plotted separately for individual mice.

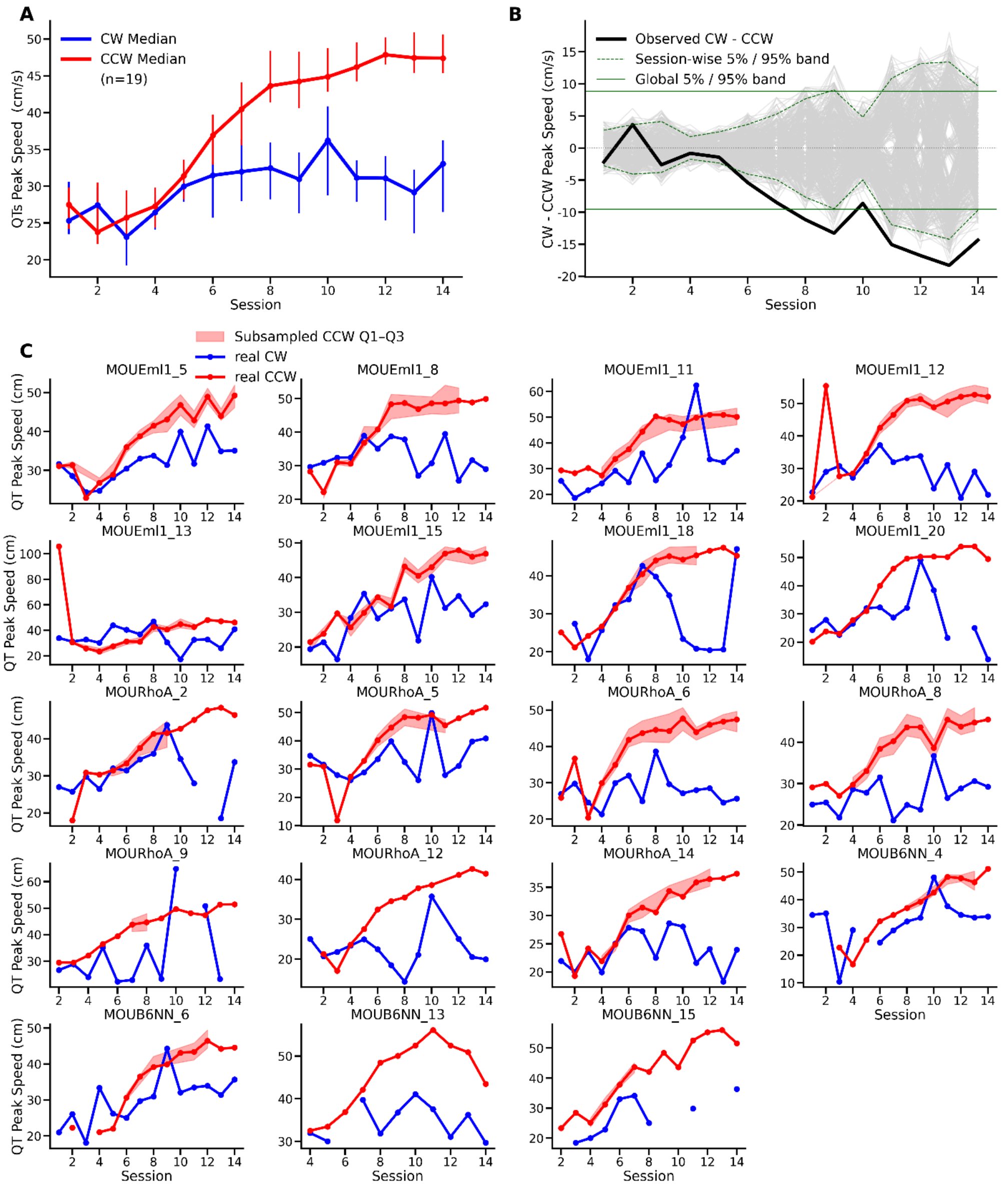


**Figure S2. Comparison of CCW and CW** QTs **peak speed across sessions. A)** Evolution of median peak speed between QTs, computed separately for CCW (rewarded direction, red) and CW (unrewarded direction, blue) turns. **B)** Permutation test comparing peak speed of CW and CCW QTs (n=500 iterations). The solid black line shows the observed difference (CW minus CCW) in median peak speed for each session. Dashed green lines indicate the global 5% and 95% confidence bands computed from the null distribution obtained by shuffling trajectory labels within each session. Solid dark green lines mark the session-wise thresholds corresponding to a z-score of ±2. Observed differences falling outside the dashed green confidence band indicate statistically significant divergence between CW and CCW QTs similarity over time. **C)** Same analysis as in panel A, plotted separately for individual mice.

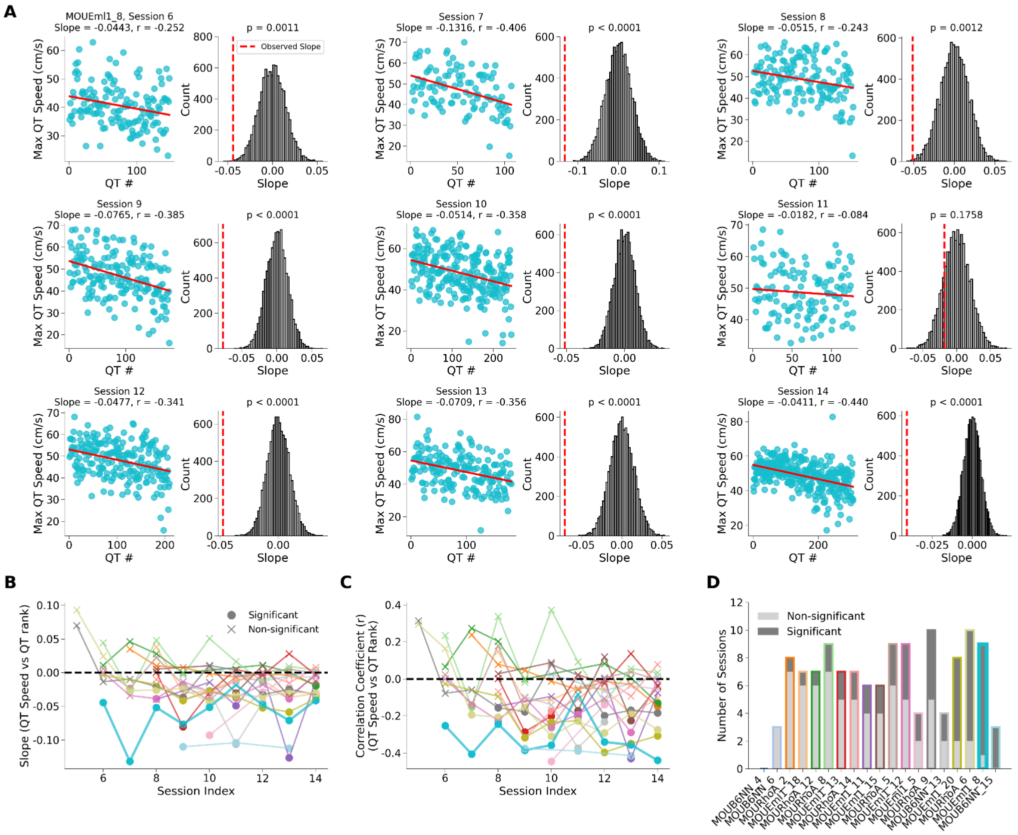


**Figure S3. Within-session changes in CCW turn speed. A)** For each of 9 example sessions, scatterplots show the maximal speed of individual CCW QTs events across their within-session trial rank. Red lines represent the linear regression fit; associated slope and correlation (r) are indicated. Right-hand panels show the corresponding null distributions of slopes from permutation tests (gray histograms), with red dashed lines indicating the observed slope and p-values reflecting the proportion of permuted slopes ≤ observed. **B)** Evolution of slope values across sessions for each mouse. Dots indicate sessions where the slope was significantly lower than expected under the null hypothesis ($p<0.05$, circle) or not (cross). **C)** Same as B, but for Pearson correlation coefficient between trial rank and CCW turn speed. **D)** p. Bar plots summarizing, for each mouse, the number of sessions with or without a significant negative slope ($p < 0.05$). Colors match individual animals across panels. Note that correlations were computed only when there were at least 100 CCW QTs, which explains why the number of sessions varies across animals.

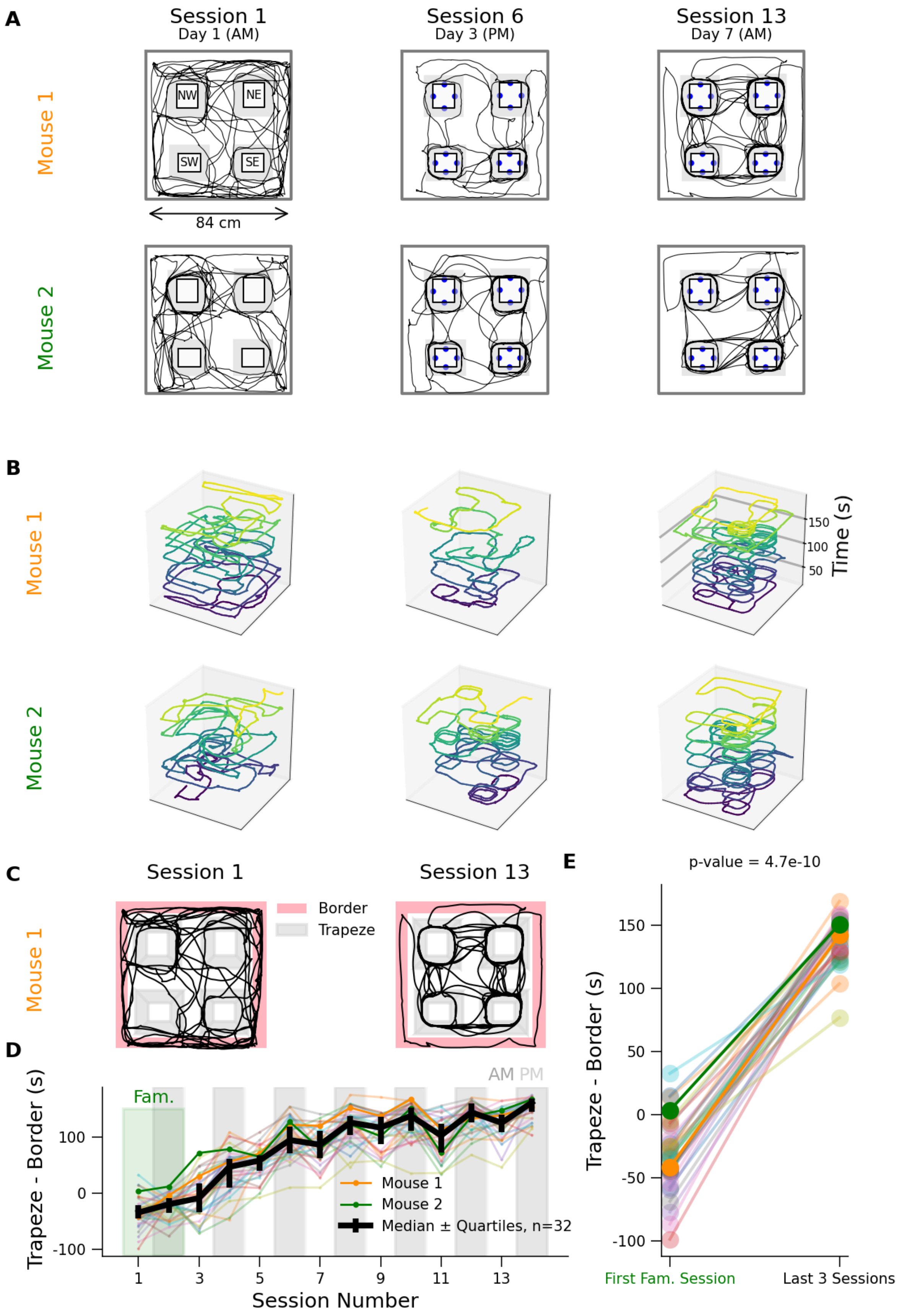
A
Session 1
Day 1 (AM)
Session 6
Day 3 (PM)
Session 13
Day 7 (AM)
Mouse 1
Mouse 2
NW
NE
SW
SE
84 cm
B
Mouse 1
Mouse 2
150
100
50
Time (s)
C
Session 1
Session 13
Mouse 1
Border
Trapeze
D
AM PM
Fam.
Trapeze - Border (s)
100
0
-100
1
3
5
7
9
11
13
Session Number
Mouse 1
Mouse 2
Median ± Quartiles, n=32
E
p-value = 4.7e-10
Trapeze - Border (s)
150
100
50
0
-50
-100
First Fam. Session
Last 3 Sessions

**Figure S4. Evolution of group 2 mice trajectories and arena occupancy across sessions.** **A**) 2D trajectories of two mice performing the 4-Towers task. Trajectories are shown for sessions 1, 6, and 13. **B**) 2D trajectories with time extended on the z-axis during the first 3 minutes of the same sessions and animals as in A, color-coded according to the session time from early (dark blue) to late (yellow). **C**) Comparison of time spent in the border and trapeze areas for Mouse 1 during 3 minutes of sessions 1 and 13. **D**) Evolution of the trapezes-border time difference across sessions. Black line represents the median and quartiles. Dark (light) grey background shows morning (afternoon) sessions, and green background shows familiarization sessions. **E**) Comparison of trapezes-border time difference between the first familiarization session and the last three sessions. Example Mouse 1 (Mouse 2) is represented in orange (green).

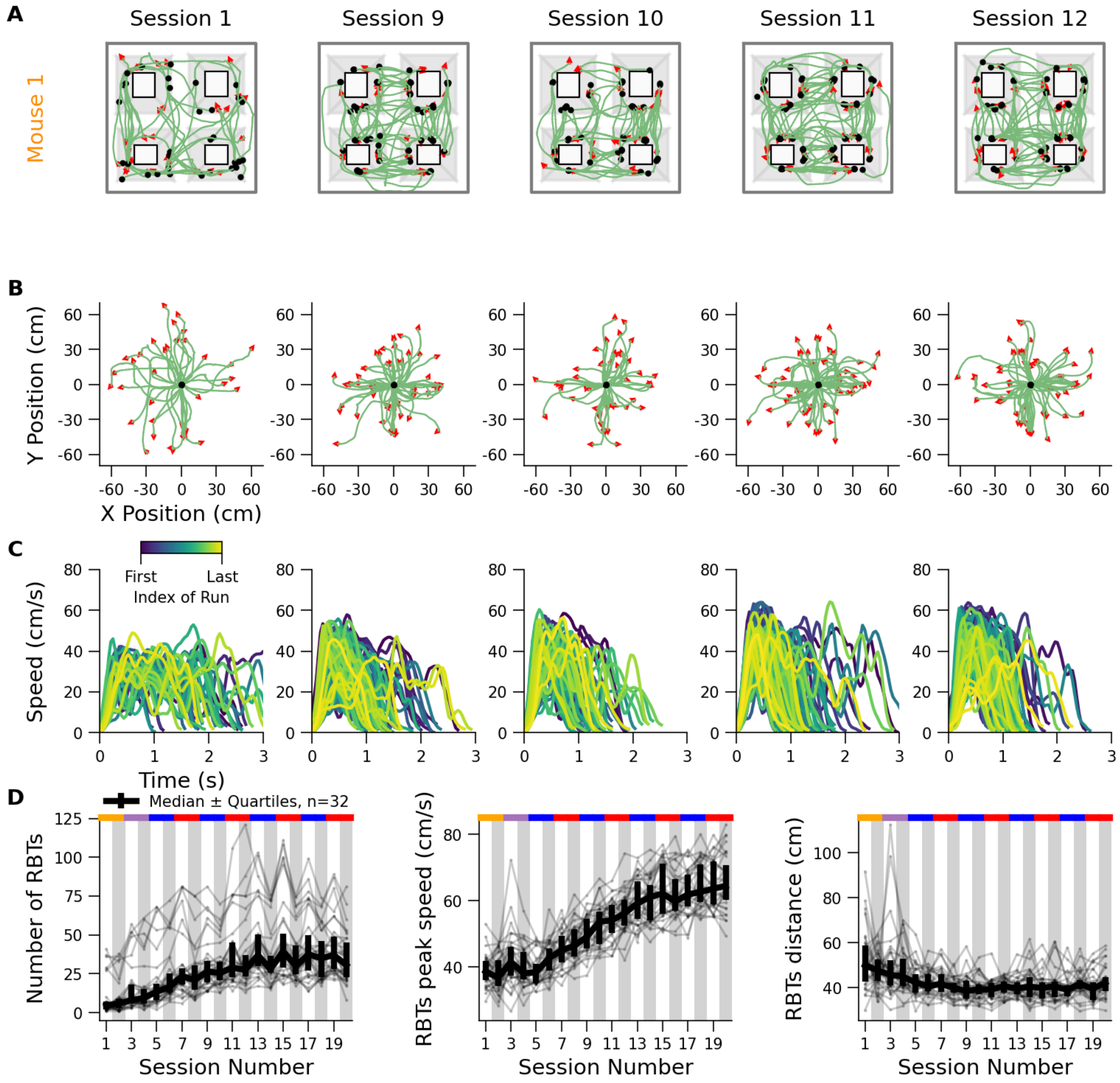


**Figure S5. Emergence of fast and short runs between towers for group 2 mice. A)** Detected RBTs of Mouse 1 for sessions 1, 7, and 14. **B)** Same as A but RBTs share the same origin. **C)** Speed profiles of RBTs for the same sessions as above, color-coded according to their rank during the sessions from early (dark blue) to late (yellow). **D)** Across sessions evolution of the number of RBTs (left), their median peak speed (center) and the median RBT distance length covered (right).

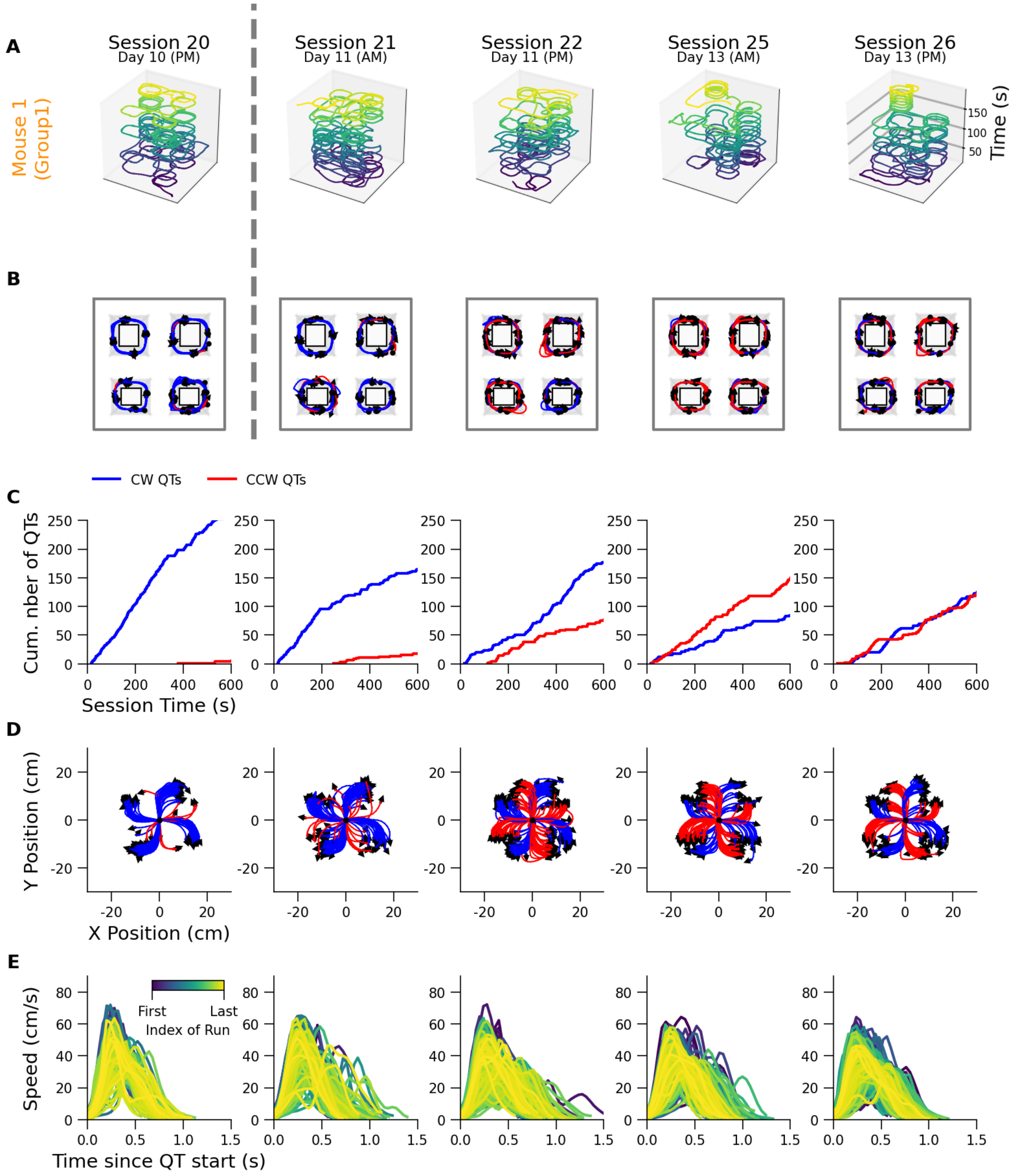


**Figure S6. Learning and adaptation of a single mouse from group 1 when the protocol is uncertain. A**) 2D trajectories with time extended on z-axis for 5 sessions (first 3 minutes; session 20 before change of protocol ; sessions 21 and 22 at the beginning of the procedure; sessions 25 and 26 at the end) for a single mouse. **B)** Detected QTs for the same mouse and sessions as A. **C)** Same as B but QTs share the same origin. **D)** Speed profiles of QTs during the same sessions, color-coded according to their rank during the sessions, from early (dark blue) to late (yellow). **E)** Cumulated number of CW and CCW QTs during the same sessions.

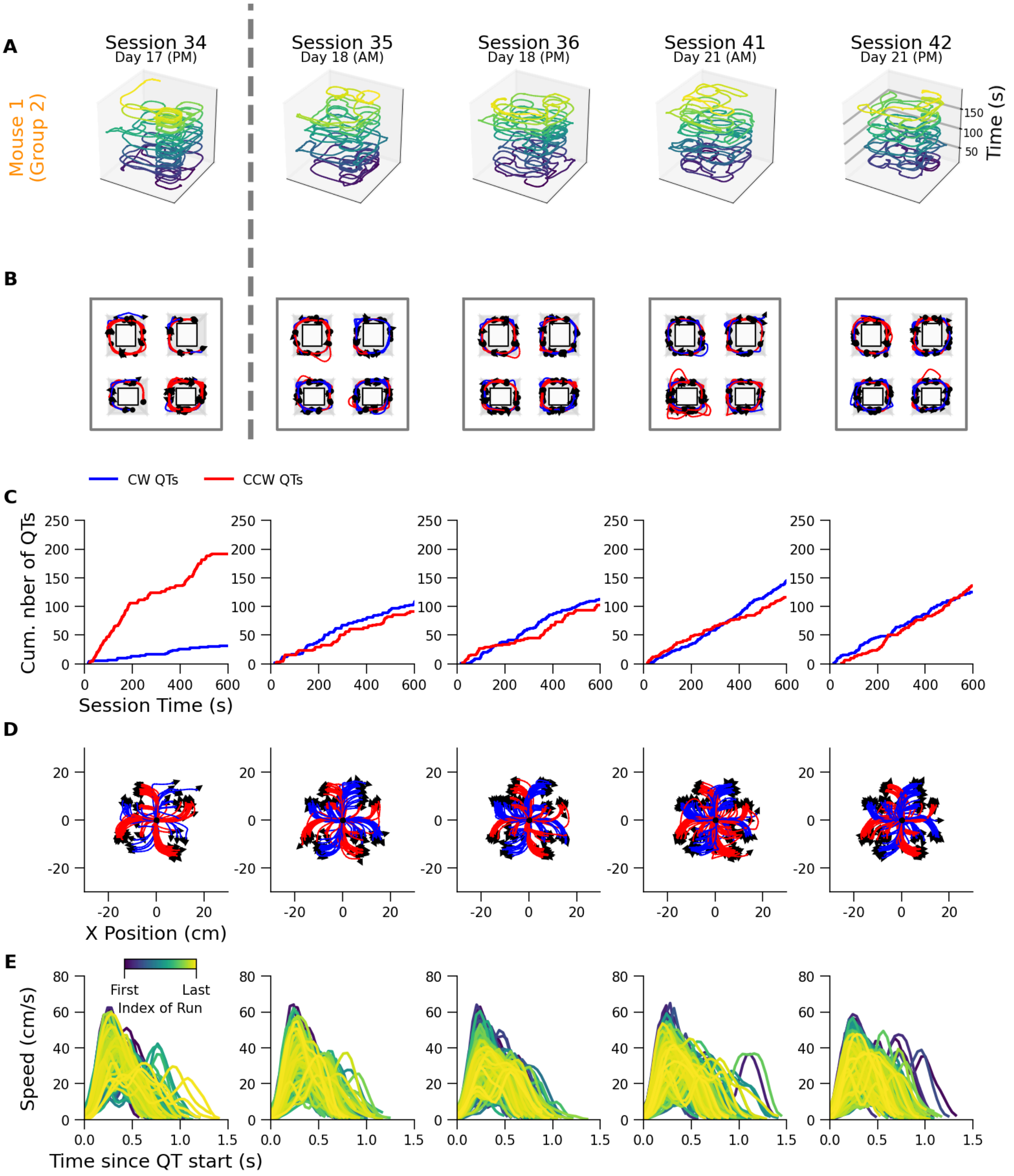


**Figure S7. Learning and adaptation of a single mouse from group 2 when the protocol is uncertain. A**) 2D trajectories with time extended on z-axis for 5 sessions (first 3 minutes; session 34 before change of protocol ; sessions 35 and 36 at the beginning of the procedure ; sessions 41 and 42 at the end) for a single mouse. **B)** Detected QTs for the same mouse and sessions as A. **C)** Same as B but QTs share the same origin. **D)** Speed profiles of QTs during the same sessions, color-coded according to their rank during the sessions, from early (dark blue) to late (yellow). **E)** Cumulated number of CW and CCW QTs during the same sessions.

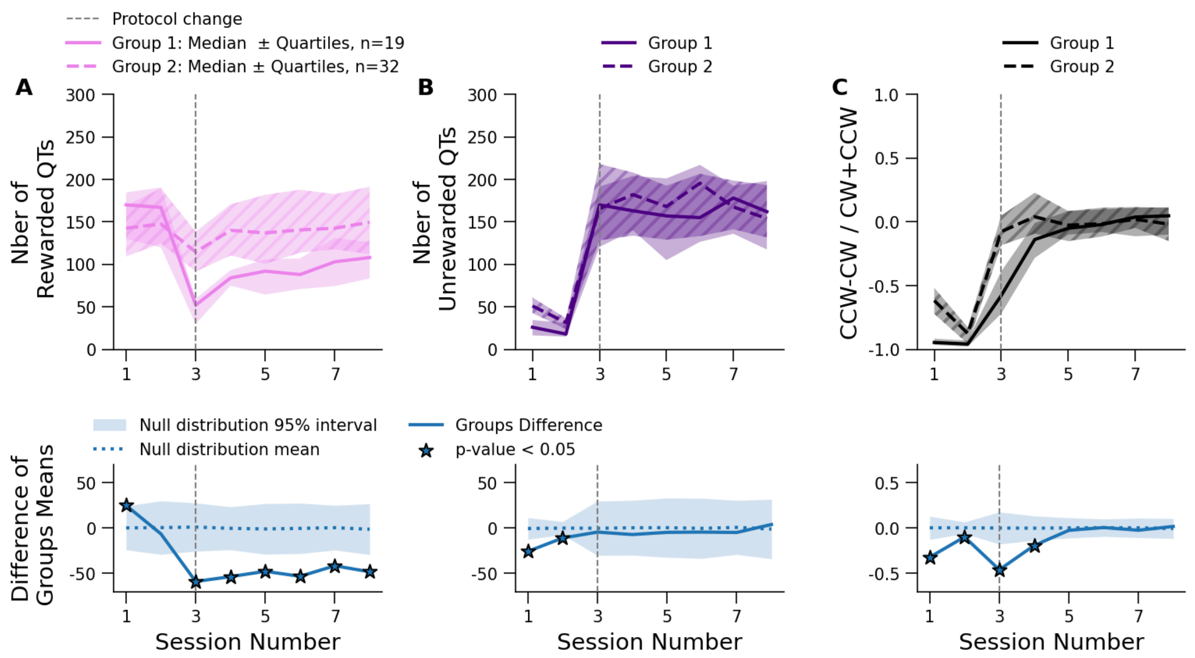


**Figure S8. Mice exposed to daily rule reversals outperform those trained under stable contingencies in a protocol requiring a high level of flexibility. A)** *Top,* number of rewarded QTs across two sessions before and six sessions after entry into the highly uncertain protocol, for the two groups (Group 1: same animals as in Figures 2–5; Group 2: same animals as in Figures 6–7). *Bottom,* difference between the two groups' mean, tested against permuted group assignments (permutation test, n=500 iterations). **B)** Same as A for unrewarded QTs. **C)** Same as A for normalized directional bias.

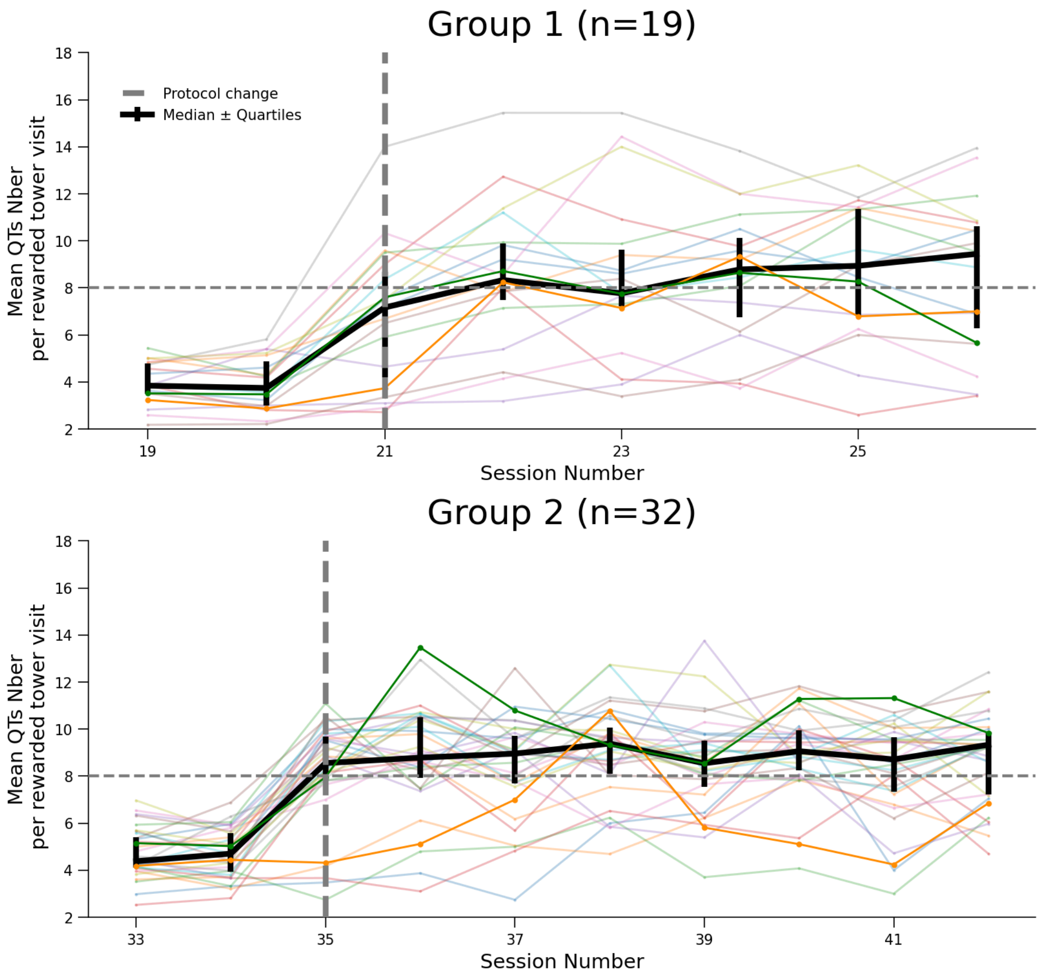


**Figure S9. Increasing perseverance in exploitation during rewarded visits of group 1 and 2 during uncertain protocol.** Mean number of QTs per rewarded visit (i.e., visits with at least one rewarded QT) across sessions for all the mice of group 1 (top, n=19) and 2 (bottom n=32).

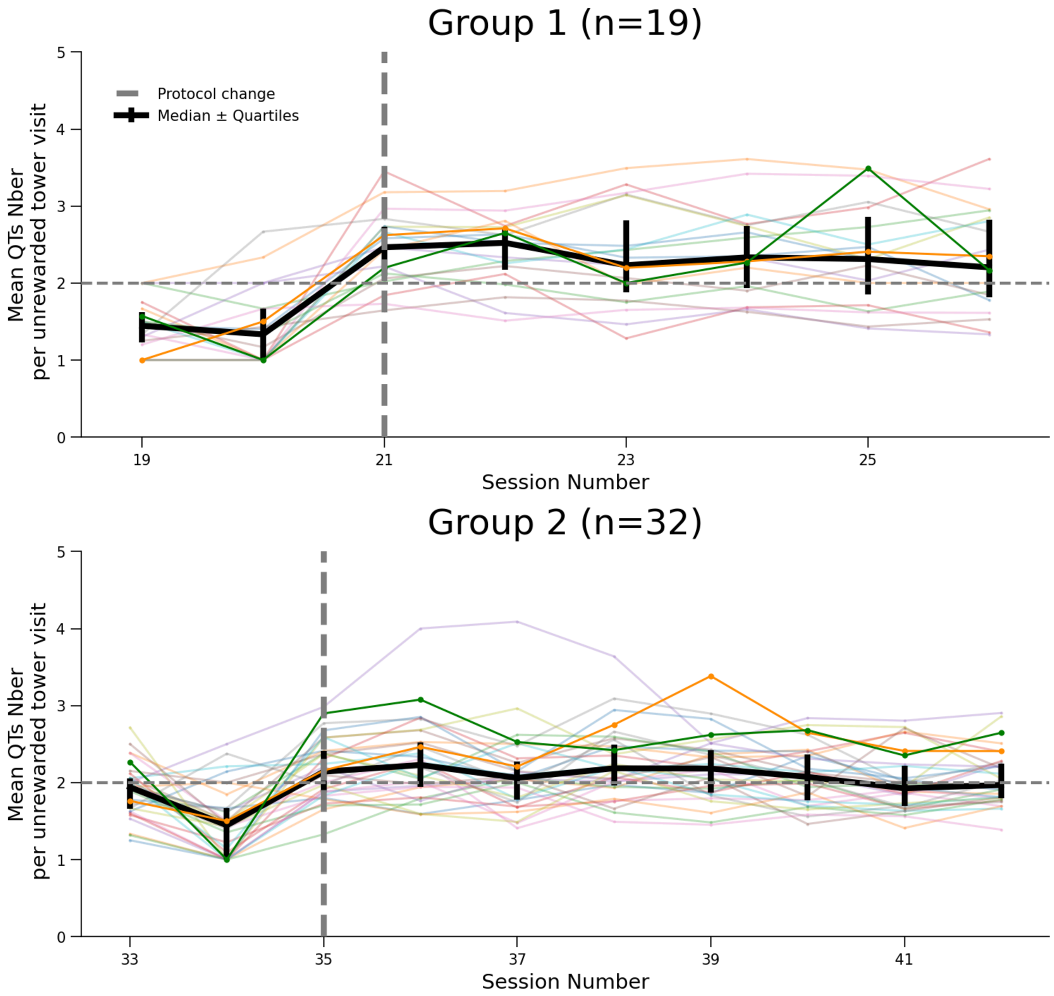


**Figure S10. Exploitation perseverance during unrewarded visits of group 1 and 2 during uncertain protocol.** Mean number of QTs per unrewarded visit (i.e., visits with no rewarded QT) across sessions for all the mice of group 1 (top, n=19) and 2 (bottom n=32).

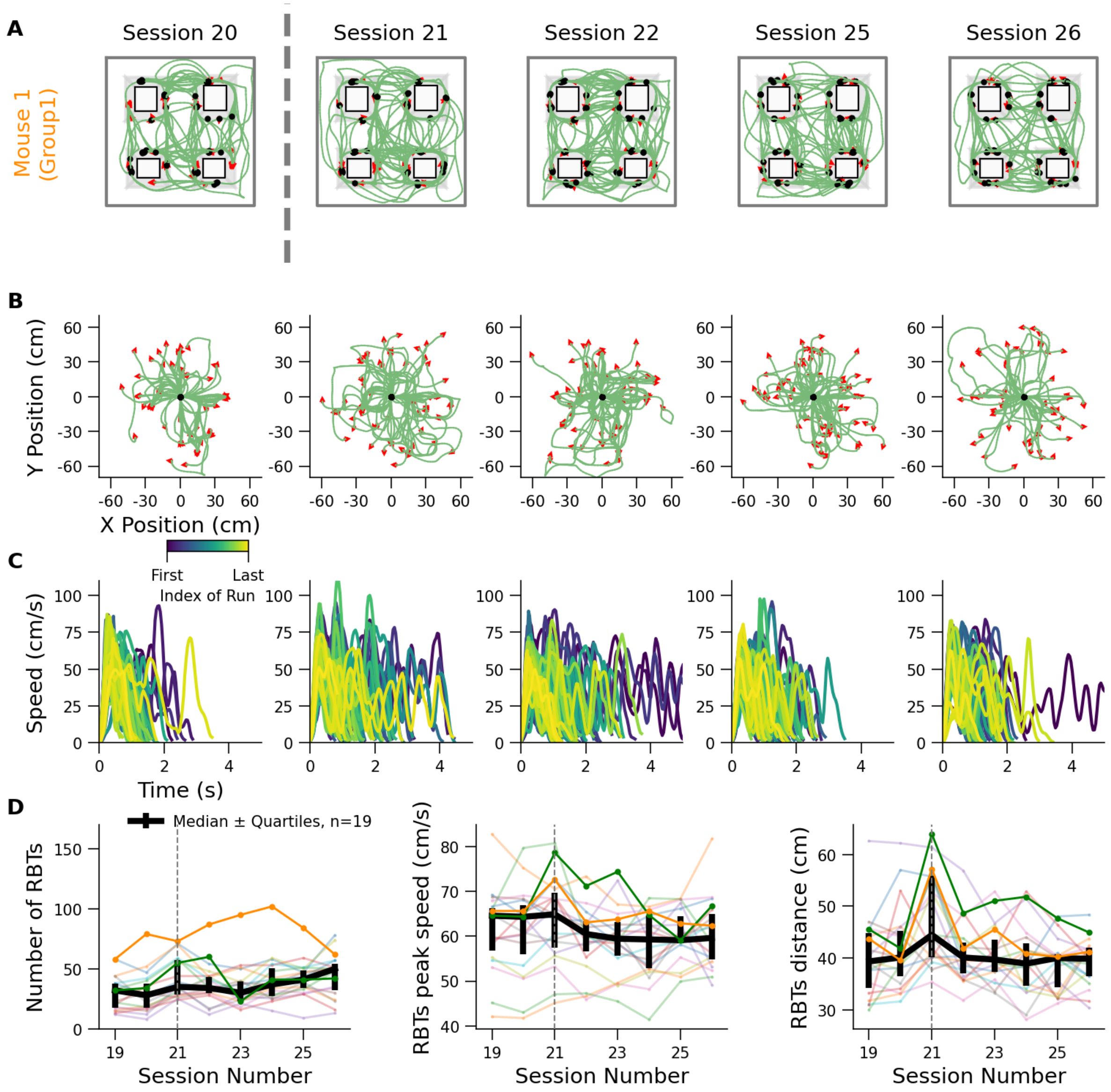


**Figure S11. Evolution of run between towers during uncertain protocol in group 1 mice. A)** Detected RBTs of Mouse 1 from group 1 for sessions 20, 21, 22, 25 and 26. **B)** Same as A but RBTs share the same origin. **C)** Speed profiles of RBTs for the same sessions as above, color-coded according to their rank during the sessions from early (dark blue) to late (yellow). **D)** Across sessions evolution of the number of RBTs (left), their median peak speed (center) and the median RBT distance length covered (right).

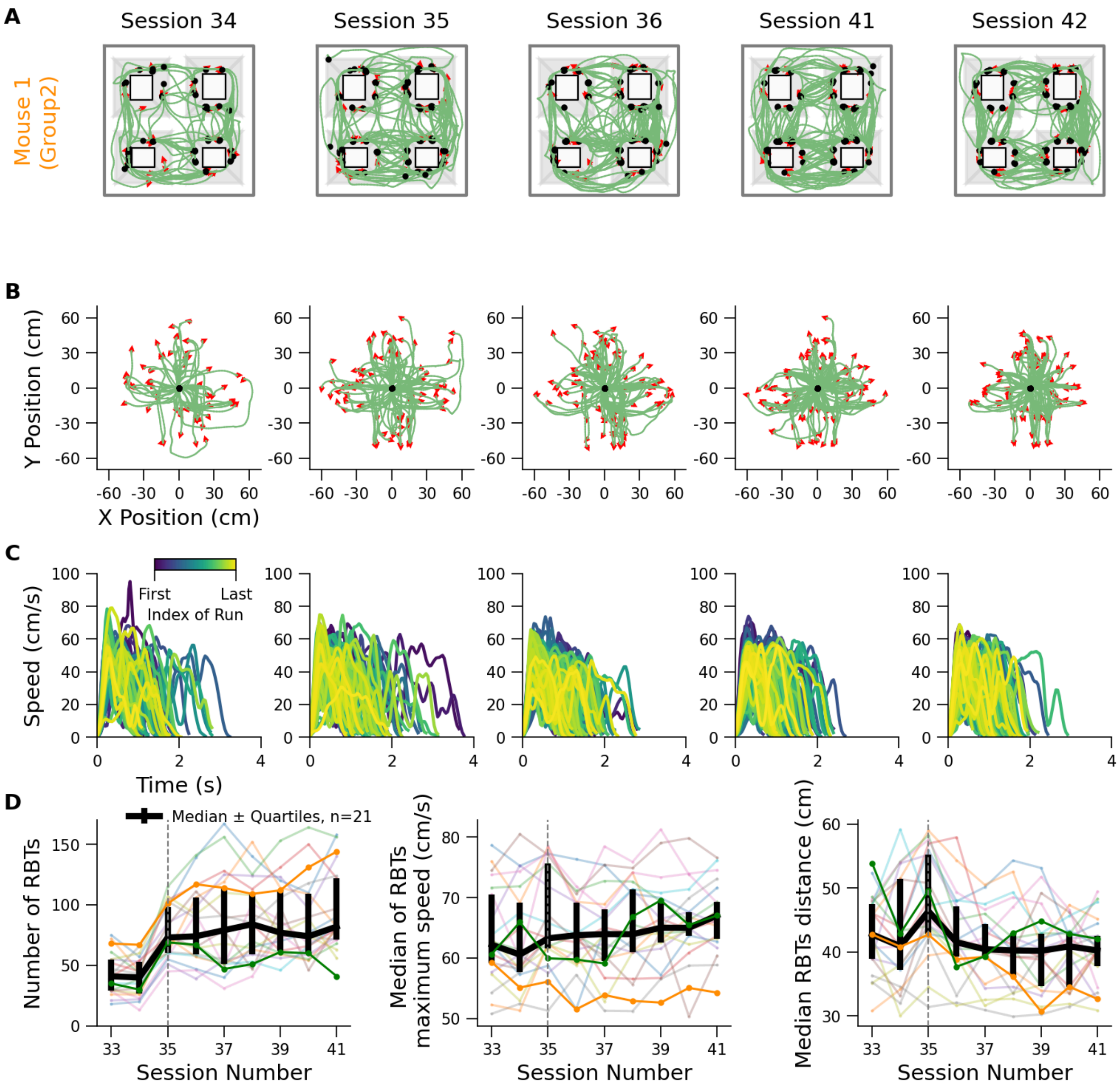


**Figure S12. Evolution of run between towers during uncertain protocol in group 2 mice. A)** Detected RBTs of Mouse 1 from group 2 for sessions 34, 35, 36, 41 and 42. **B)** Same as A but RBTs share the same origin. **C)** Speed profiles of RBTs for the same sessions as above, color-coded according to their rank during the sessions from early (dark blue) to late (yellow). **D)** Across sessions evolution of the number of RBTs (left), their median peak speed (center) and the median RBT distance length covered (right).
